# Abrogation of presynaptic facilitation at hippocampal mossy fiber synapses impacts neural ensemble activity and spatial memory

**DOI:** 10.1101/2024.09.10.612312

**Authors:** Catherine Marneffe, Noelle Grosjean, Kyrian Nicolay-Kritter, Cécile Chatras, Evan R. Harrell, Ashley Kees, Christophe Mulle

**Affiliations:** Interdisciplinary Institute for Neuroscience, CNRS UMR 5297, University of Bordeaux, F-33000 Bordeaux, France

**Keywords:** presynaptic plasticity, synaptotagmin 7, memory, hippocampus, CA3, mossy fiber synapses

## Abstract

Presynaptic short-term plasticity is thought to play a major role in the process of spike transfer within local circuits. Mossy fiber synapses between the axons of dentate gyrus (DG) granule cells and CA3 pyramidal cells (Mf-CA3 synapses) display a remarkable extent of presynaptic plasticity which endows these synaptic connections with detonator properties. The pattern of action potential firing, in the form of high frequency bursts in the DG, strongly controls the amplitude of synaptic responses and information transfer to CA3. Here we have investigated the role of presynaptic facilitation at Mf-CA3 synapses in the operation of CA3 circuits *in vivo* and in memory encoding. Syt7, a calcium sensor necessary for presynaptic facilitation, was selectively abrogated, in DG granule cells using Syt7 conditional KO mice (DG Syt7 KO mice). In hippocampal slices, we extend previous analysis to show that short-term presynaptic facilitation is selectively suppressed at Mf-CA3 synapses in the absence of Syt7, without any impact on basal synaptic properties and long-term potentiation. Short-term plasticity was found to be crucial for spike transfer between the DG and CA3 in conditions of naturalistic patterns of presynaptic firing. At the network level, in awake head-fixed mice, the abrogation of short-term plasticity largely reduced the co-activity of CA3 pyramidal cells. Finally, whereas DG Syt7 KO mice are not impaired in behavioral tasks based on pattern separation, they show deficits in spatial memory tasks which rely on the process of pattern completion. These results shed new light on the role of the detonator properties of DG-CA3 synapses, and give important insights into how this key synaptic feature translate at the population and behavioral levels.

## Introduction

It is widely accepted that storage and retrieval of information depend on multiple forms of activity-dependent synaptic and intrinsic plasticity processes with distinct temporal dynamics (Takeuchi et al., 2014). In this context, most of the attention has focused on long-term changes in synaptic efficacy driven mostly by changes in the number and/or conformation of postsynaptic receptors and only modify synaptic gain. However, neurons often encode information not as spikes in isolation but as bursts of 2–20 spikes at high frequency (>20Hz), which triggers a wide range of presynaptic forms of synaptic plasticity (short- and long-term) depending on the synaptic contacts and on the history of synaptic activity (Jackman & Regehr, 2017; Monday et al., 2018). Presynaptic plasticity mostly relies on changes in neurotransmitter release probability, which in turn alter the short-term dynamics of synapses. Short-term synaptic plasticity (STP) modulates the efficacy of synaptic transmission on timescales from a few milliseconds to several minutes (Zucker & Regehr, 2002), and as such has a direct influence on network-level dynamics which take place on the time scale of stimulus-driven activity and behavior (Tsodyks & Markram, 1997).

Synaptic facilitation is a form of STP which occurs at synapses with a low basal probability of release, at which an isolated action potential often fails to elicit any postsynaptic response and to reliably evoke action potentials in the postsynaptic neuron. At such weak synapses, facilitation can dramatically increase release probability, especially following high frequency bursts of incoming action potentials (Fuhrmann et al., 2002; Lisman, 1997; Rotman et al., 2011). Computational modelling of facilitating synapses predicts they are optimized for information transfer at presynaptic rates of 9-70 Hz, with less than 10 incoming action potentials (Fuhrmann et al., 2002). Recently, progress has been made in understanding the mechanisms of presynaptic facilitation. It is thought that synaptic facilitation corresponds to an increase in vesicle release as a consequence of a larger calcium transient available for the second of two closely spaced stimuli (Jackman & Regehr, 2017). Activity-dependent increase in calcium entry into the synaptic terminal has been shown to play a role through presynaptic spike broadening or facilitation of voltage-gated calcium currents (Jackman & Regehr, 2017). Synaptotagmin7 (Syt7) acts as the major Ca^2+^-sensor binding residual Ca^2+^ to increase release probability and promote synaptic facilitation (Jackman et al., 2016). Constitutive genetic deletion of Syt7 was shown to disrupt synaptic facilitation at excitatory synapses in the hippocampus and thalamus (Jackman et al., 2016).

Short-term synaptic dynamics greatly varies between neuronal connections (Zucker and Regehr, 2002). Amongst facilitating synapses in the brain, hippocampal mossy fiber (Mf) synapses connecting the dentate gyrus to CA3 pyramidal cells (CA3-PCs, Mf-CA3 synapses) show a remarkably high dynamic range of presynaptic plasticity (Rebola et al., 2017; Vandael & Jonas, 2024) (Marneffe et al, 2024 accepted for publication). Three main forms of short-term facilitation are distinguished at Mf-CA3 synapses (Rebola et al., 2017; Vandael & Jonas, 2024) (Marneffe et al, accepted for publication): 1) Frequency facilitation, which includes low frequency facilitation and train facilitation, operating in the range of tens of milliseconds to several seconds (Salin et al., 1996); 2) Post-tetanic potentiation triggered by trains of high frequency stimulation, which lasts several minutes (Griffith, 1990; Langdon et al., 1995); 3) Finally, potentiation of synaptic transmission induced by retrograde signaling after the firing of post-synaptic CA3 PCs with an onset and offset of several minutes. By virtue of STP, Mf-CA3 synapses act as “conditional detonators” discharging their postsynaptic CA3 PC targets with bursts of presynaptic activity from single Mfs (Henze et al., 2002; Sachidhanandam et al., 2009), or with patterns of spiking readily observed in DG cells of awake mice (Chamberlain et al., 2013; Sachidhanandam et al., 2009; Vandael et al., 2020).

Even though Mf–CA3 PC synapses are highly regulated on a second to minute timescale through presynaptic short-term plasticity, the role of presynaptic facilitation in information processing in DG- CA3 circuits and in memory remains largely elusive. The discovery of Syt7 as a calcium sensor selectively involved in synaptic facilitation makes it possible to clarify the functional roles of facilitation at identified synapses, in neuronal computation and related behaviors. STP induced by short trains of presynaptic action potentials at Mf-CA3 synapses is abrogated in constitutive Syt7 KO mice. Here we have taken advantage of conditional Syt7 KO (Syt7 cKO) mice, to further characterize the impact of DG- selective deletion of Syt7 on presynaptic plasticity and information transfer between the DG and CA3, in slices and *in vivo*. In addition, we have evaluated the consequences of impaired STP at Mf-CA3 synapses on hippocampus-dependent behavioral tasks.

## METHODS

### Animals and viral transduction

All experiments complied to the European Union guidelines and were performed under the authorization of the ‘Pole in Vivo’ of the IINS following the regulations from the Ethical Committee of Bordeaux (CEAA50). The Syt^em4(IMPC)H^ mice were obtained from the MRC Harwell Institute which distributes these mice on behalf of the European Mouse Mutant Archive. Males and females were used for this study. They were housed with littermates and isolated only for surgery recovery and during the behavioral tests. They were injected (P25-P35) under isoflurane anaesthesia with pLenti225-C1ql2- ChIEFTom-2a-VSKCre (LV-Cre) or pLenti-C1ql2-2A-ChIEFTom (LV-Control) (Barthet et al., 2018) injected stereotaxically at AP -2.2; ML +/-2.0 and DV-2.3 in the hippocampus. The viruses were produced by TransBioMed, the dedicated facility at the University of Bordeaux. To maximize the infection, 1 µl was injected in each hemisphere at 100 nl/min (World Precision Instruments); titres were at least 1.13 E+09 particles/ml for LV-Cre and 6.37 E+08 particles/ml for LV-Control. The mice were used for experiments after a minimum of 20 days following viral transduction.

### *Ex vivo* slice electrophysiology

Mice (P50-P70) were anesthetized with a mix of ketamine (100 mg/kg, IP) and xylazine (10 mg/kg, IP). Mice were intra-cardially perfused with an ice-cold oxygenated cutting solution (87 mM NaCl, 2.5 mM KCl, 1.25 mM NaH2PO4, 25 mM NaHCO3, 25 mM glucose, 75 mM sucrose, 0.5 mM CaCl2, 7 mM MgCl2, pH 7.4, 315 mOsm/kg). The brain was rapidly removed, 300 µm thick parasagittal sections were obtained with a vibratome (Leica, VT1200S, Germany), which were then incubated in a resting chamber containing ACSF (125 mM NaCl, 2.5 mM KCl, 1.25 mM NaH2PO4, 26 mM NaHCO3, 1.3 mM MgCl2, 2.3 mM CaCl2, 10 mM glucose, pH 7.4, 300 mOsm/kg) for 20 minutes at 34°C. After recovery, the slices were kept at room temperature, in oxygenated (95% O2, 5% CO2) ACSF.

Borosilicate glass capillaries (Harvard apparatus) were pulled (Sutter Instruments) to obtain 3-6 MΩ patch pipettes. For voltage clamp experiments, the patch pipette was filled with an intracellular solution containing: 140 mM Cs-methanesulfonate, 2 mM MgCl2, 4 mM NaCl, 5 mM Na-phosphocreatine, 0.2 mM EGTA, 10 mM HEPES, 3 mM Na2-ATP, 0.3 mM GTP, P, pH 7.2 adjusted with CsOH, 280 mOsm/Kg. The series resistance was assessed by a voltage step (-10 mV, 5 ms) at the beginning of each recorded sweep, and the recording was discarded if the series resistance varied over 20 MΩ or by more than 20%. No correction for the liquid junction potential was applied. The data was recorded with an EPC 10 amplifier (HEKA Elektronik), filtered in Patch Master software (Lambrecht, Germany) with a Bessel filter at 2.9 kHz, digitized at 10 kHz.

The recordings of evoked CA3 excitatory post synaptic currents (EPSCs) were performed in the voltage clamp mode at room temperature in presence of 10 µM bicuculline to inhibit GABA-A receptors. The stimulation was performed by 1 ms light pulses over the DG *stratum moleculare* and the baseline was recorded at 0.1 Hz stimulation. Train facilitation was recorded with 10 consecutive pulses at 20 Hz, evoked at an interval of 50 ms. The inhibitory post synaptic currents (IPSCs) were recorded at room temperature in presence of 50 µM DAP-V at + 5 mV, the reversal membrane potential for cations. The same protocols of light stimulation were used as for the EPCS.

The recordings of fEPSPs (field excitatory post synaptic potentials) in slices were performed with a patch pipette filled with ACSF. After a 20 minute recording for baseline acquisition, 3 sweeps of 125 light pulses at 25 Hz were applied. At the end of the recording, LCCG-1 was applied to verify that the stimulation effectively triggered Mf-CA3 synaptic responses.

### *In vivo* awake head-fixed silicon probe recordings

At P30, the mice were unilaterally injected with LV-Cre or LV-Control in the DG and implanted with a head bar to allow for later head fixation. The head bar was fixed to the skull with dental cement. To avoid covering with cement the region above the hippocampus and thereby preventing later inserting of the probe to perform the recordings, a plastic ring was placed on the skull, above the region covering DG and CA3 and filled with removable silicon. 7 days after the implantation, the mice were head-restrained in a stereotaxic frame over a wheel and was allowed to freely explore for habituation for 10 minutes. The next days, the mice were head fixed over the wheel for increasing periods of time: 10, 20, 40 and then 60 minutes. The mice were then trained daily for 60 minutes for 20 days after the head bar implantation. This period of time was selected to allow for the lentivirus to express the construct and to maximize the training on the wheel. On the day of the recording, the mice were anesthetized and the silicon was removed to have access to the skull. In a ∼30 minutes surgery, two craniotomies were performed: one over the injection site onto the DG and a second bigger one antero-laterally to access CA3. The brain surface was constantly maintained humid with a saline solution. The mice were allowed to recover for an hour and were then head fixed for the recording. One probe was inserted in the DG (Neuronexus A1x32-6mm-50-177) and connected to a 32-channel adaptor (Neuronexus, Adpt- A32-OM32), and the other in CA3 (A4x16-poly2-5mm-23s-200-177) connected to a 64-channel adaptor (Neuronexus, *Adpt-A64-OM32x2-sm*). The adaptors were connected to Intan headstages (Intan RHD 32 #C3314 and RD 64 #C3315). The wheel systems were provided by Polywheel (Imetronic, France). The probes were inserted in the brain at a speed not exceeding 1 µm/second using a micromanipulator (Scientifica). The assessment of the location was performed by constant monitoring of depth and of the brain signal. When both probes were positioned in the expected location, we started a recording session of 30 minutes. At the end, the probes were taken out and the mice were euthanized for post-hoc brain histology and verification of probe location and virus expression.

### Quantification and statistical analysis of *in vivo* electrophysiological data

*Local Field Potential.* Raw (.dat) data files for LFP and membrane potential were read into Python using the Neo package (Garcia et al., 2014), and signal processing was performed with custom routines using the SciPy package (Virtanen et al., 2020). LFP signal was downsampled 16 times to 1250 Hz and filtered using a 4-pole Butterworth band-pass filter between 0.2 Hz and 300 Hz.

*Spike Sorting.* The spike sorting was performed with Spyking CIRCUS (Yger et al., 2018) and hand-curated in Phy (https://github.com/kwikteam/phy). The unit classification for excitatory and inhibitory cells was done as in Stark et al., 2013 (Stark et al., 2013), with an added condition that putative pyramidal cells had to have an average firing rate lower than 5 Hz.

*Brain state classification.* Individual theta cycles were detected as in Lopes-Dos-Santos, et al., 2018, (Lopes-dos-Santos et al., 2018), using an EMD filter and finding the oscillations that matched theta frequency (4-12 Hz). Theta cycles that occurred within 250 ms of one another were joined into theta epochs. Only theta epochs that were longer than 550 ms were included in subsequent analyses. Any time the wheel speed was greater than 0 was counted as running. Periods when there was running and/or theta were considered the ‘active’ state, and all other times were considered ‘inactive’.

*SWR and dentate spike detection.* In our dataset 5 Ctl and DG-Syt7 KO mice also had good recordings in CA1. CA1 sharp wave ripples were detected with the CNN model described in Navas-Olive et al. 2022, (Navas-Olive et al., 2022). In our dataset, 4 Ctl and 5 DG-Syt7 KO mice displayed dentate spikes in the LFP of the dentate gyrus. We detected them using the python packaged shared by the Dupret lab (https://github.com/mcastelli98/DentateSpikeClassifier).

*Population Activity.* Two measures of population activity in CA3 were used: pairwise correlation coefficients, and population activity events. Pairwise correlation coefficients were found by first making spike trains by binning the firing of CA3 PCs into 20 ms bins. Then, for each pair of neurons, the correlation coefficient of the spike trains was taken. A higher correlation coefficient indicates that a particular pair of cells tend to fire within the same 20 ms time bins. The second measure of CA3 population activity used was the population event. These were found by first making the spike trains in 20 ms bins for all the CA3 PCs, as before. Then, in each bin, the number of cells active was counted. Then we calculated a threshold (in number of neurons active) for each session. We did this by shuffling the spike trains from that session and finding the number of simultaneously active neurons that occurred less than 5% of the time. We considered this the threshold for that session because it represented the number of simultaneously active cells that was unlikely to occur by chance. Time bins where the activity exceeded this threshold were considered population events.

*Statistics.* Parametric and nonparametric statistics were performed with the SciPy package (Virtanen et al., 2020) for Python. Bootstrap statistics were performed with custom Python routines. Unless otherwise noted, data were expressed as the median +/- the mean absolute derivation about the median, which was calculated by subtracting each data point from the group’s median, taking the absolute value, summating, and then dividing by the number of data points. These measures of central tendency and variation were used because many distributions were found not to be normal. When comparing 2 independent groups, for example, when comparing Ctl and DG-Syt7 KO groups, a Mann-Whitney U test was performed. A bootstrap test was used when comparing two non-independent groups, for example, for comparing between the active and nonactive states within one animal. In this test, the difference between paired data points was randomly determined to be either positive or negative 1,000 times. If the mean of the differences in the actual data set was more extreme than 95% of those in the shuffled data set, the paired groups were considered to have a significant difference.

### Histology

After the *in vivo* recordings, the mice were deeply anesthetized with isoflurane, dislocated and their brain was quickly taken out of the skull. For the assessment of Syt7 levels in unilaterally injected animals, the mice were injected with a mix of ketamine (100 mg/kg, IP) and xylazine (10 mg/kg, IP) then intra-cardially perfused with 50 ml PBS, followed by PFA 4% until complete stiffness. For all mice, after an overnight fixation in PFA 4%, the brains were transferred in PBS. 50 µm coronal slices were acquired with a vibratome (Leica, VT1200S, Germany). Slices were permeabilized at room temperature in 0.3% Triton X100 for two hours, then incubated overnight at 4°C with Syt7 (Synaptic systems 105173, 1/250) primary antibody. After rinsing in PBS, the slices were incubated with secondary antibody (ThermoFisher, A11008, 1/1000) at room temperature for 2 hours. The slices were mounted with Dapi-containing mounting solution (Vectashield, H-1200-10) and imaged with a Nanozoomer (Hamamatsu Photonics).

### Pattern separation tasks

We used two separate tasks to test for the ability of the mice for pattern separation, which is the ability to discriminate representations of two overlapping memories or similar experiences (Cès et al., 2018). We first used a spatial pattern separation task. The two first days, the animals went through a habituation phase: the first day, mice were placed for ten minutes in an 1 X 1 m open field without any objet and the second day, mice were habituated to the presence of 2 two random objects. The test phase occurred during three days: the mice were placed in the field with two test objects during 12 minutes (sample phase). After a 30-minutes inter-trial period, the mice were placed back in the arena for 6 minutes with a fixed object in the same position as during the sample phase, and the second object was moved 20, 35 or 45 cm away from the first object. The mice were presented all the distances in random order during 3 days.

We then used an object pattern separation task. During the sample phase, the mice were placed in a 45 x 45 cm arena with two objects for 10 minutes. After an inter-trial period of 4 minutes, one of the objects was replaced with another one expressing highly similarity or low similarity with reference to the replaced object. Mice were tested for both low and high similarity objects in a random order during 2 days. For both tests, the calculated interaction ratio was measured as the interaction time of the changed (or moved object) divided by the total interaction time with the two objects. Data were only included if the mouse interacted with the objects during the test phase.

### Pattern completion task

We designed a task in a Barnes maze to test for pattern completion, the ability to retrieve a memory from a degraded or partial cue input. Based on the mouse natural avoidance for open enlighten spaces, the Barnes maze is used to assess their ability to remember an escape box location relying on a full set of cues, a partial set of cues or in the absence of cues. The apparatus consists in a 90 cm diameter table with 18 holes of 38 mm diameters. Under one of the holes, an escape box of 13 cm by 9 cm is fixed. Around the table, walls had 15 visual cues. First, mice were habituated to the escape box by placing them in the box during 60 sec and then were allowed to explore during 5 min. During the acquisition training phase (three days, 6 trials per day), the mouse was placed for each trial in the center of the maze in a opaque cylinder during around 15 sec, the cylinder was lifted and the mouse was allowed to explore during 2 min. If the mouse did not find the escape box, it was gently guided to it. The time and the distance to reach the box was measured. The last trial of the last day was counted as a probe with all cues. On day 4, only a partial set of cues was presented and the box was removed. The mouse was allowed to explore during 2 min (Probe “partial cues”). On day 5 none of the cues were left on the walls and the mouse was allowed to explore during 2 min without the box (Probe “minimal cues”). The latency to first reach the box was measured for all probe tests. The behavior of mice was recorded with a camera (Noldus). The distance they walked and time to find the escape box was measured with a software (EthoVision XT 15 by Noldus Information Technology). The statistical analyses were performed with Graph Pad Prism.

### Anxiety assessment

To assess anxiety, we first analyzed the mouse locomotion and trajectories in an open field. All behavioral recordings were filmed with a camera (Noldus) and the data were processed with EthoVision XT 15 by Noldus Information Technology. We then placed the mice in an elevated plus maze (EPM) with two enclosed arms and two open arms and tracked their avoidance of the open arms. The mice were allowed to freely explore the apparatus for 10 minutes. Finally, we monitored the burying behavior of the mice in the marble burying task. A cage was prepared with 20 marble beads placed at the surface of a 10 cm layer of litter. A picture was taken at the time that the mouse was placed in the cage. After 15 minutes, the mouse was taken out of the cage and a second picture taken. The burying ratio was then calculated based on the remaining visible surface of beads on top of the litter over the initial surface.

## RESULTS

### Adult onset DG-specific deletion of Syt7 combined with optogenetic stimulation

By injecting Syt7 floxed mice with a lentiviral construct expressing ChIEF, Tomato and Cre under a C1ql2 promoter (LV-Cre), we selectively expressed Cre recombinase in DG granule cells leading to the deletion of the Syt7 gene and to a combined expression of channel rhodopsin to allow for light stimulation (Fig. 1A,(Barthet et al., 2018)). The LV-Cre and LV-Control (same construct except it lacks Cre) constructs express Tomato as a reporter gene to allow for rapid screening of infection in the hippocampus. In mice infected with LV-Cre (hereafter named DG-Syt7 KO), the expression of ChIEF was restricted to cells expressing Cre, which allowed Mf-EPSCs to be recorded from a population of neurons lacking Syt7.

**Figure 1.**
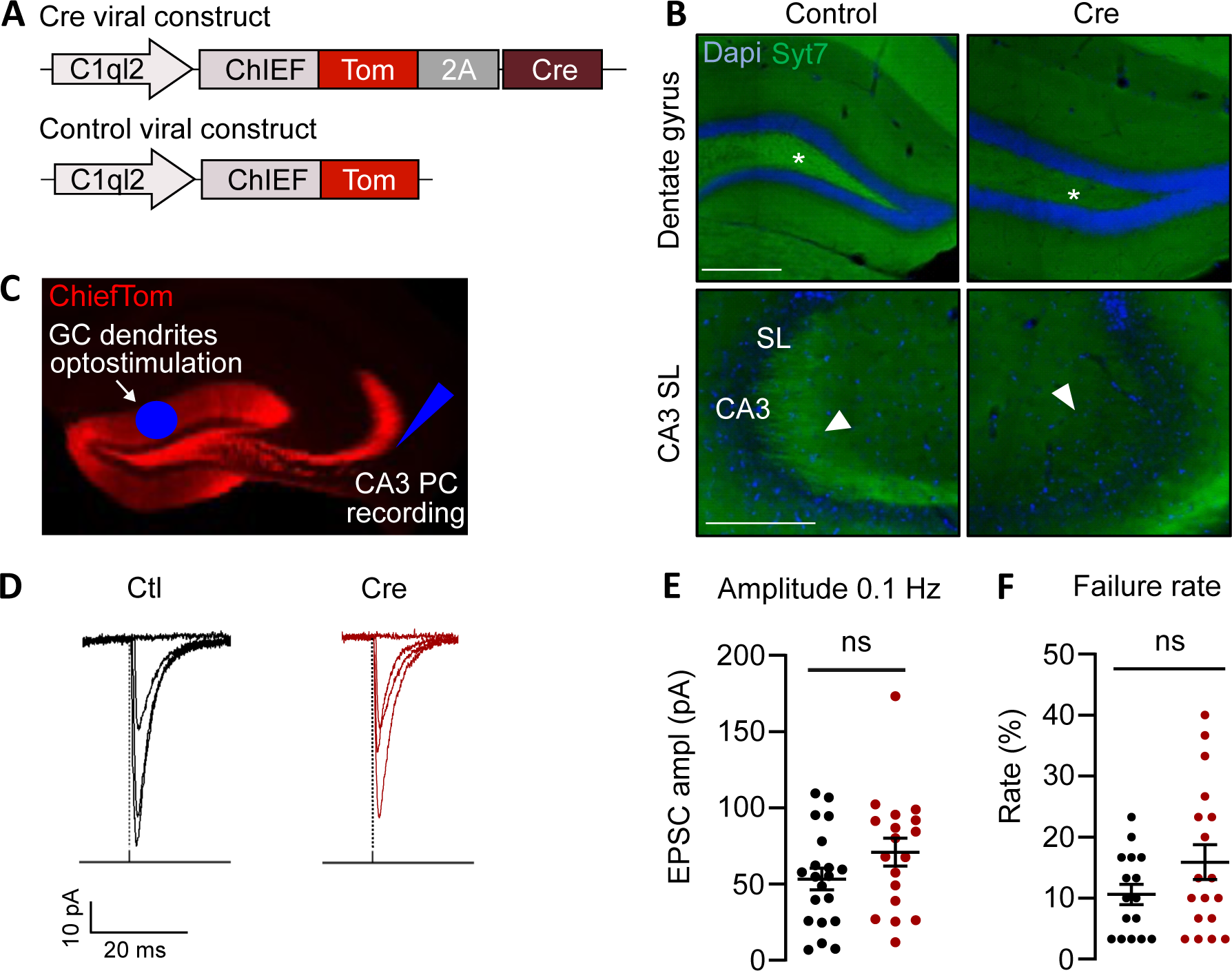
Selective removal of Syt7 in presynaptic mossy fiber terminals by expression of LV-C1ql2- ChIEF-Tom-2A-Cre in the dorsal DG does not change basal properties of MF transmission. **A**. Syt7Flox mice are injected with a lentivirus expressing Cre under a DG-specific promoter. Scheme of the used lentivirus: C1ql2 promoter drives the expression of ChIEF, Tomato, a 2A self-cleaving peptide, and the Cre recombinase. The control virus is the same construction except that the Cre sequence is omitted: pLenti225-C1ql2-ChIEFTom-2a-VSKCre (LV-Cre) or pLenti-C1ql2-2A-ChIEFTom9/10/24 8:24:00 PM (LV-Control). **B**. Syt7 expression in DG-Syt7 KO mice (here and throughout the figures indicated as Cre mice) is strongly decreased in the hilus and in *stratum lucidum*. Scale bar 200 μm. **C**. Patch clamp recordings in slices are performed in CA3 PCs, and Mf-CA3 EPSCs are evoked by pulses of blue light in the DG molecular layer. Bicuculline is added to the bath to block GABAA receptor activity. Illus- tration of the optical stimulation in the DG dendritic field and recording in CA3 PC: expression of Tomato (Tom) is restricted to the dendrites of DG granule cells and to the mossy fiber (Mf) track. **D**. Example traces of Mf-CA3 EPSCs evoked by light stimulation at 0.1 Hz in Ctl and DG-Syt7 KO (Cre) mice. **E.** No difference in basal synaptic transmission properties is observed at Mf-CA3 synapses with 0.1 Hz stimulation between the two groups. Scatter plot of the mean +/- SEM are presented for recorded amplitudes at 0.1 Hz stimulation (n = 24 in Ctl, mean ampl = 53.3 pA; n = 22 in DG-Syt7 KO, mean ampl = 70.9 pA, Mann-Whitney, p = 0.1579). **F.** Failure rates are unchanged for the two groups (Ctl = 10.6%, DG-Syt7 KO = 15.9%, p = 0.2784).

We first confirmed the efficiency of infection with LV-Cre in deleting Syt7 by performing immunostaining of Syt7 in unilaterally injected mice (Fig. 1B). Syt7 is ubiquitously expressed in the brain, and is enriched in the hilus and in the *stratum lucidum* (SL) where Mfs make synaptic contacts with CA3 PCs (Barthet et al., 2018), in line with its role in synaptic facilitation at Mf-CA3 synapses (Jackman et al., 2016). We measured the mean pixel intensity of Syt7 staining in the *stratum lucidum* and normalized it to the mean pixel intensity in *stratum lucidum* in the contralateral side. We observed an average decrease of Syt7 of 59.6% stats (n= 4, Mann-Whitney, p= 0.0286) in the *stratum lucidum*, providing an estimation of the percentage of DG cells transduced with the LV-Cre construct. Of note, although the injection targeted the dorsal hippocampus, the ventral hippocampus was also in large part infected by the LV-Cre viral construct (Sup Fig. 1).

### Removal of Syt7 from DG cells abrogates presynaptic facilitation in Mf-CA3 synapses

We bilaterally injected LV-Cre or LV-Control in Syt7^fl/fl^ mice and recorded Mf-CA3 EPSCs by performing patch-clamp recordings of CA3 PCs and optogenetic stimulation of granule cell dendrites in the DG (Fig. 1C, D). As previously shown (Barthet et al., 2018), focal light stimulation (1 ms) of the somatodendritic region of the DG at 0.1 Hz evoked EPSCs with amplitudes and failure rates similar to those evoked by minimal electrical stimulation (Marchal & Mulle, 2004), strongly suggesting that the recorded Mf-CA3 EPSCs corresponded to the stimulation of single Mfs. This allowed to obtain a faithful comparison of Mf-CA3 EPSC properties in WT vs. DG-Syt7 KO synapses (Fig. 1E, F). No difference was observed in the average amplitude at 0.1 Hz (n = 24 in Ctl, mean ampl = 53.2 pA; n = 22 in DG-Syt7 KO, mean ampl = 70.9 pA, Mann-Whitney, p = 0.1579) or the failure rate (Ctl = 10.6%, DG-Syt7 KO = 15.9%, p = 0.2784), strongly suggesting that the absence of presynaptic Syt7 did not impair initial release probability nor number of active release sites. We observed that the amplitude and frequency of spontaneous EPSCs was not different between the genotypes (Sup Fig. 2). We also found that the proportion of EPSCs with amplitudes >40 pA, which likely represent EPSCs of Mf origin (Henze et al., 1997), were not significantly changed, indicating that Syt7 removal does not affect spontaneous release at Mf-CA3 synapses.

**Figure 2.**
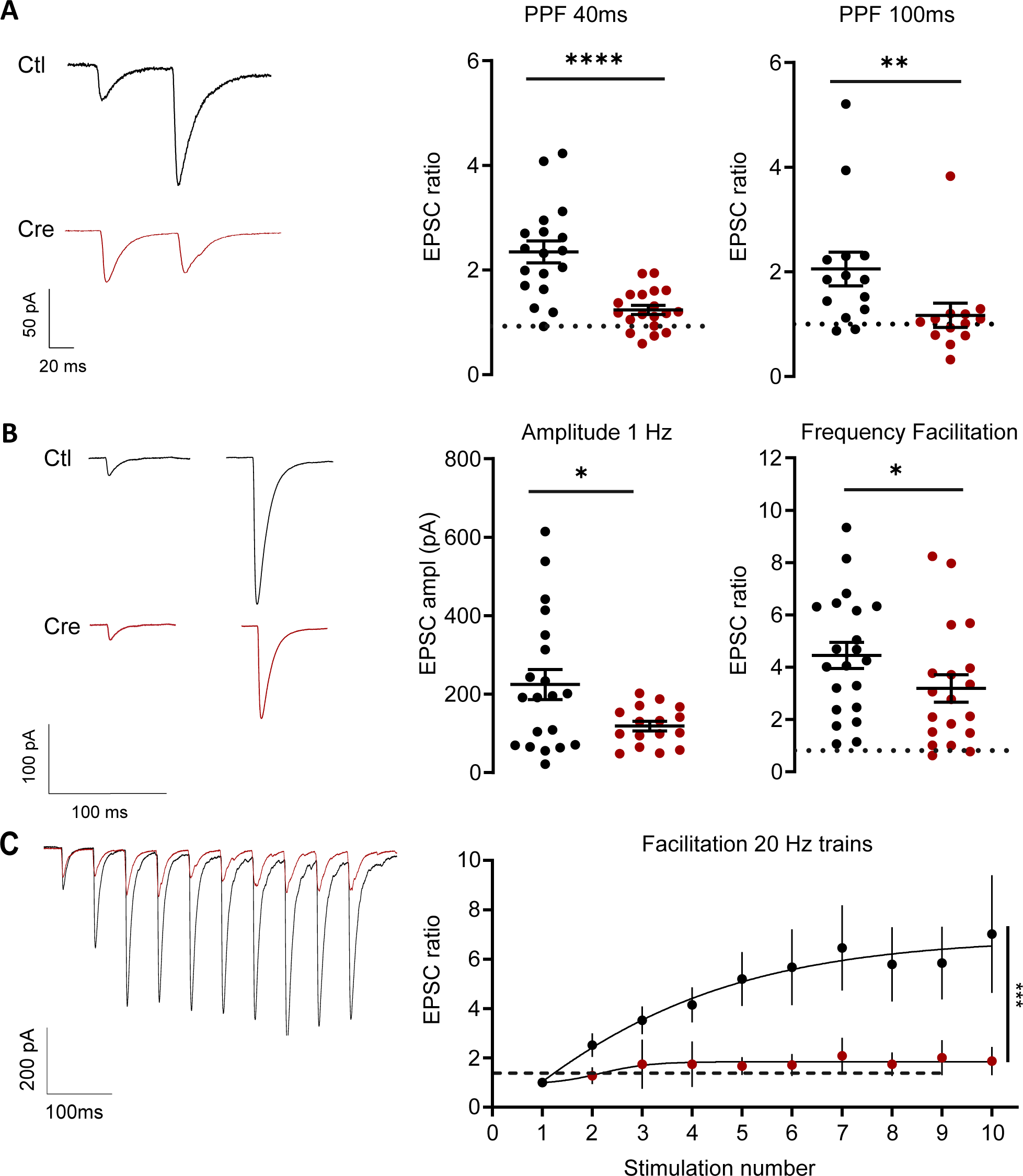
Synapse specific abrogation of presynaptic short-term plasticity at Mf-CA3 synapses. **A**. Left: average traces for paired-pulse stimulation at 40 ms interval in Ctl and DG-Syt7 KO (Cre) mice. Paired-pulse facilitation is calculated as the ratio of the amplitude of the second EPSC amplitude over the first (Ctl = 2.37, DG-Syt7 KO = 1.24, Mann-Whitney, p<0.0001). Right: quantification of paired-pulse stimulation at 100 ms interval (Ctl = 1.85, DG-Syt7 KO = 1.04, p =0.0032). **B**. Representative traces of light-evoked Mf-CA3 EPSCs at 1 Hz in Ctl and DG-Syt7 KO (Cre) mice. The amplitude of Mf-CA3 EPSCs at 1 Hz is decreased in DG-Syt7 KO mice (1 Hz Ctl = 233.7 pA, Cre = 98.9 pA, unpaired t-test, p = 0.0227) due to a decrease of low frequency facilitation (ratio of the amplitude at 1 Hz vs. 0.1 Hz) (Ctl = 4.25, DG-Syt7 KO = 2.77, Mann-Whitney, p = 0.0472). **C**. Superimposed example traces of presynaptic facilitation in 2 response to 20 Hz trains in Ctl (black) and DG-Syt7 KO (red) mice. Facilitation is measured as the ratio of the amplitude of Mf-CA3 EPSCs for each light pulse normalized to the amplitude of the first Mf-CA3 EPSC in the train. Presynaptic facilitation is strongly diminished in DG- Syt7 KO mice (Ctl, n=13, DG-Syt7 KO, n = 15, two-way Anova, p = 0.0074).

When applying paired-pulse stimulation at either 40 ms or 100 ms intervals of stimulation, a robust decrease in the EPSC ratio was observed at DG-Syt7 KO vs. Ctl Mf-CA3 synapses (Fig 2A, 40 ms: p<0.0001; 100 ms: p = 0.0032, Mann-Whitney, n = 14 in Ctl and n = 13 in DG-Syt7 KO mice). We observed a moderate decrease in low-frequency facilitation when changing the rate of stimulation from 0.1 Hz to 1 Hz (p = 0.0472, Mann-Whitney, n = 14 in Ctl and n = 13 in DG-Syt7 KO mice) (Fig. 2B). Finally, the removal of Syt7 from Mf-CA3 synapses lead to an almost total loss of presynaptic facilitation in response to short trains of 10 pulses of light at 20 Hz (Fig. 2C, n = 13 in Ctl, n = 15 in Cre, two-way Anova, p = 0.0074). Overall our findings with selective abrogation of presynaptic Syt7 at Mf- CA3 synapses confirm the observation made in constitutive Syt7 full KO mice using fEPSP recordings (Jackman et al., 2016). In addition, our results also indicate that the basal synaptic properties (probability of release at 0.1 Hz, number of release site and quantal size) are not affected by the selective abrogation of presynaptic Syt7 at Mf-CA3 synapses.

### Presynaptic PTP and LTP are not impaired at DG-Syt7 KO Mf-CA3 synapses

Mf-CA3 synapses display a prominent form of presynaptic long-term potentiation (LTP), independent of the activation of NMDA receptors, which is classically induced by a series of high frequency trains (100 Hz)(Nicoll & Schmitz, 2005). Presynaptic LTP can also be induced by optogenetic stimulation of Mfs (3 times 125 light stimulation at 25 Hz) (Monday et al., 2022). To check if Syt7 was involved in presynaptic LTP, we recorded fEPSPs in the *stratum lucidum* while optically stimulating Mfs in the hilus. We recorded a 20-minute baseline of fEPSPs evoked at a rate of 0.1 Hz before applying light pulses 3 x 125 times at 25 Hz (Fig 3 A-C). The extent of post-tetanic potentiation (PTP), measured as the average fEPSP facilitation during 1 minute following the induction protocol, was not significantly different between Ctl and DG-Syt7 KO Mf-CA3 synapses (Fig 3D, Ctl = 529 %, n = 13 slices; DG-Syt7 KO = 473 % n = 14 slices, Mann-Whitney, p = 0.3527). The extent of LTP was averaged during a 10 minute period 60-70 minutes following LTP induction. No significant difference was observed in the magnitude of LTP between control and DG-Syt7 KO Mf-CA3 synapses (Fig. 3E, Ctl = 274 %, n = 13 slices; DG-Syt7 KO = 230 %, n = 14 slices, Mann-Whitney, p = 0.2588). This result indicates that a high level of short-term presynaptic facilitation is dispensable for presynaptic LTP at Mf-CA3 synapses. Overall, the DG-Syt7 KO mice allow selective assessment of the consequences of abrogated short-term facilitation (at a ms to seconde time scale) at Mf-CA3 synapses, without any change in basal synaptic properties, in PTP or in LTP, on neural circuit mechanisms and related behavior.

**Figure 3.**
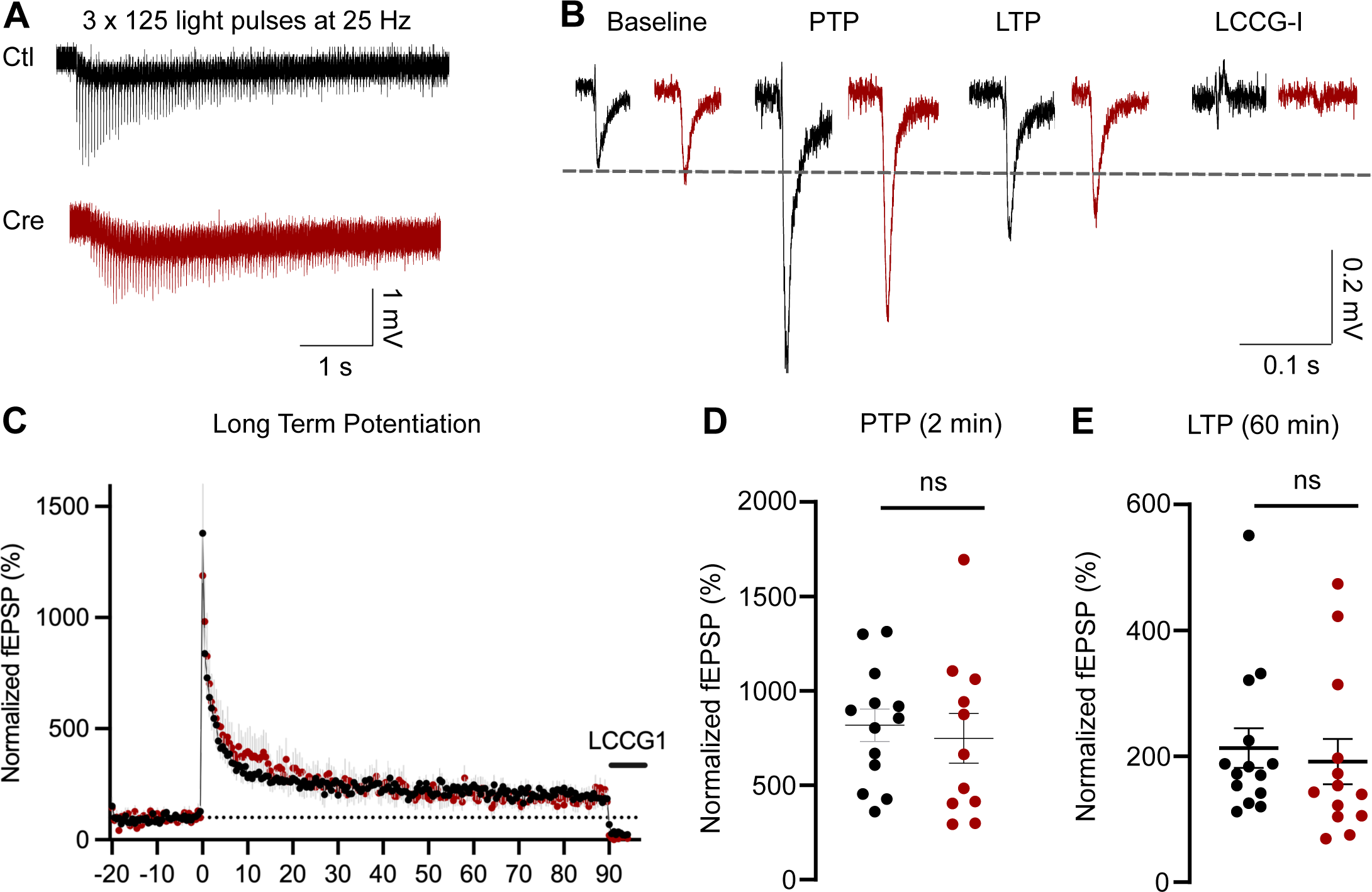
Selective removal of Syt7 in DG granule cells does not abrogate PTP and presynaptic LTP at Mf-CA3 synapses. **A**. Example traces of a train of a train of 125 light pulses at 25 Hz used to trigger PTP and LTP in control (black) and in DG-Syt7 KO (Cre, red) mice. An extracellular electrode was positioned in the stratum lucidum to record fEPSPs. **B**. Example traces of fEPSPs triggered by the light pulse at a baseline 0.05 Hz stimulation (10 minutes before the induction train stimulations), post-tetanic potentiation (2 minutes after the trains), LTP (40 minutes after trains), and LCCG-I (10 minutes after drug application). **C**. Time course of the amplitude of fEPSP represented +/- SEM for control (black) and DG-Syt7 KO mice (red).**D.** Average values recorded from 1 minute after train stimulation (Ctl = 854.9, DG-Syt7 KO = 663.5, Mann-Whitney, p = 0.5309. **E.** Average values recorded from 60 to 70 minutes after train stimulation (Ctl = 213.0, DG-Syt7 KO = 191.7, Mann-Whitney, p = 0.2588

### Feedforward inhibition is decreased in DG-Syt7 KO mice

We then assessed how abrogation of STP at Mf-CA3 synapses impacted spike transfer at DG-CA3 connections by first addressing feedforward inhibition in DG-Syt7 KO mice. Indeed, mossy fibers not only contact CA3 PCs but also send filopodial extensions which make synapses onto interneurons forming the anatomical basis of feedforward inhibition. To indirectly examine whether the deletion of Syt7 also affected excitatory Mf synapses onto inhibitory interneurons, we recorded IPSCs evoked in CA3 PCs by light stimulation of DG granule cells and by clamping the cell membrane at +5 mV, the reversal potential for cations. Although the amplitude of the di-synaptic IPSCs showed a large variability in basal stimulation conditions for both genotypes, we observed a significant decrease in the average IPSC amplitudes in the absence of Syt7 at Mf-CA3 synapses at both a frequency of 0.1 Hz and 1 Hz (at 0.1 Hz: p = 0.1745, at 1 Hz: p = 0.2396, Two-way Anova, Ctl, n=21; DG-Syt7 KO, n=11) (Fig. 4A,B). Similar findings were observed in train stimulations where the absolute amplitude of the inhibitory response was smaller in the DG-Syt7 KO mice, although no facilitation was observed (Fig. 4D, p= 0.0187, Two-way Anova, Ctl, n=21; DG-Syt7 KO, n=11). Feedforward inhibition appears to be generally impaired in the absence of Syt7 at the excitatory Mf-CA3 synapses, possibly as a compensatory mechanism to counteract the loss of synaptic facilitation. We measured the ratio of EPSC amplitude (at -70 mV) vs. IPSC amplitude (at +5 mV) for each stimulation during 0.1 Hz, 1 Hz, and in 20 Hz trains (Fig. 3). The ratio of amplitudes at 1 Hz vs. 0.1 Hz was not significantly affected (Fig. 4E, Fig. 4E, p= 0.1027, Two-way Anova, Ctl, n=21; DG-Syt7 KO, n=11). When normalized to the first stimulation in the 20 Hz train, we found that the E/I ratio was progressively reduced along the train of stimuli at DG-Syt7 KO Mf-CA3 synapses, mainly due to impairment of facilitation of Mf-CA3 EPSCs (Fig. 4F, p= 0.0147, Two-way Anova, Ctl, n=21; DG-Syt7 KO, n=11).

**Figure 4.**
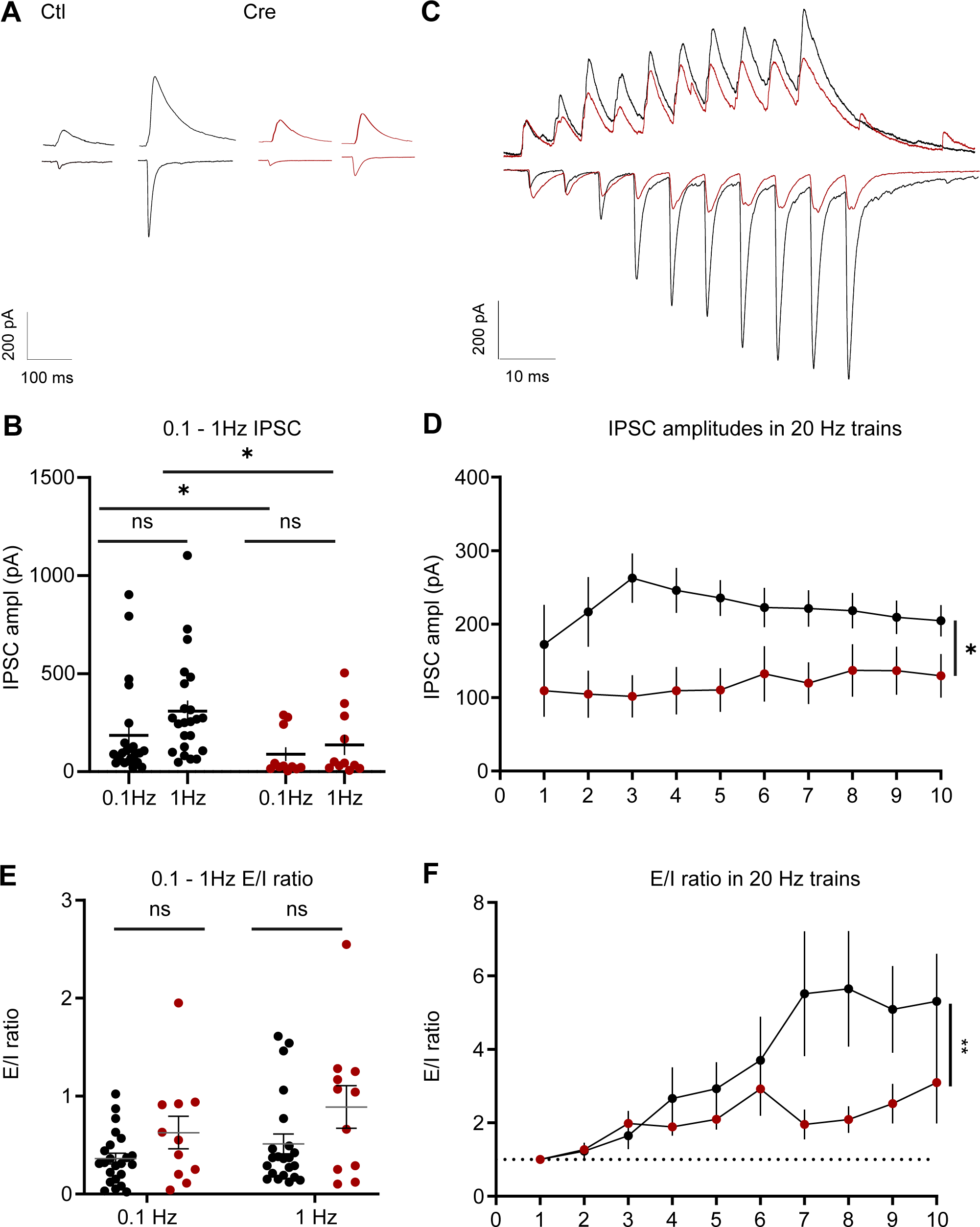
Impact of DG selective Syt7 deletion on Mf-CA3 feedforward inhibition. **A**. EPSCs are recorded at -70 mV in presence of 50 μM DAP-5 then IPSCs are recorded from the same CA3 PCs at +5 mV. Example traces of recorded IPSCs in control and DG-Syt7 KO (Cre) mice in baseline (0.1 Hz) conditions. **B**. Scatter plots of the mean IPSC amplitudes are presented, showing a difference in the average IPSC amplitude recorded at both 0.1 Hz and 1 Hz stimulation rates (0.1 Hz: Ctl = 185.2 pA, n=21; and DG-Syt7 KO = 89.7 pA, n=11; Mann-Whitney, p = 0.0230; 1 Hz: Ctl = 308.8 pA, and DG- Syt7 KO = 136.5 pA, Mann-Whitney, p = 0.120). No low frequency facilitation (from 0.1 to 1 Hz) was observed in any of the genotypes. **C**. Example traces of superimposed EPSCs and IPSCs in response to a 20 train stimulation in control (black) and DG-Syt7 KO (red) mice. **D.** Graph of the average amplitude of IPSCs during the 20 Hz stimulation train in Ctl (black) and DG-Syt7 KO (red) mice. Whereas a marked change in average IPSC amplitude is observed for all stimulations, no significant facilitation of IPSC amplitude is found along the train. **E.** Scatter plots of the mean E/I ratios calculated at the ratio of average EPSC amplitude over average IPSC amplitude for each cell recorded. A large variability in the values was observed, and a significant difference was found between Ctl (black) and DG-Syt7 KO (red) mice at low frequencies of stimulation. **F.** Graph of the E/I ratio normalized to the value obtained to the first stimulation in the train. The E/I ratio increases during the 20 Hz train likely due to facilitation of EPSCs. A large difference in the level of increase of the E/I ratio between control and DG-Syt7 KO mice is observed (p= 0.0147, Two-way Anova, Ctl, n=21; DG-Syt7 KO, n=11).

### Spike transfer at Mf-CA3 synapses of DG-Syt7 KO mice with a natural stimulus pattern

We then examined what consequences the loss of facilitation and the decreased E/I ratio may have in the transfer of spikes across Mf-CA3 synapses in response to a naturalistic train of stimulation. We used a pattern of high-frequency DG granule cell activity (recorded from awake mice - see *in vivo* recordings below), lasting 30 sec and which consisted of 57 spikes with variable inter-event intervals (IEI) resulting in instantaneous frequencies ranging from 1 Hz to 75 Hz (Fig. 5A). This natural pattern of spiking activity was used to optically stimulate DG granule cells in slices from Ctl and DG-Syt7 KO mice, while recording CA3 PCs in the current-clamp mode, in the absence of any inhibitor of receptors for neurotransmitters (Fig. 5A). As expected, actions potentials were mostly triggered in periods of high frequency repeated stimulation. The stimulus pattern was repeated 7 times with a 2 minutes interval, and spiking activity was averaged (Fig. 5B), demonstrating a robust decrease in spike transfer from the DG to CA3 PCs along the whole duration of natural stimulus pattern presentation. In Ctl mice, the spike discharge probability increased progressively as a function of instantaneous frequency of presynaptic stimulation from 1 to > 50 Hz (Fig. 5C) (p<0.0001; Two-way Anova; Ctl, n=8; DG-Syt7 KO, n=13). Overall, abrogation of short-term presynaptic facilitation at Mf-CA3 PC synapses leads to a profound impairment in spike transfer from the DG to CA3.

**Figure 5.**
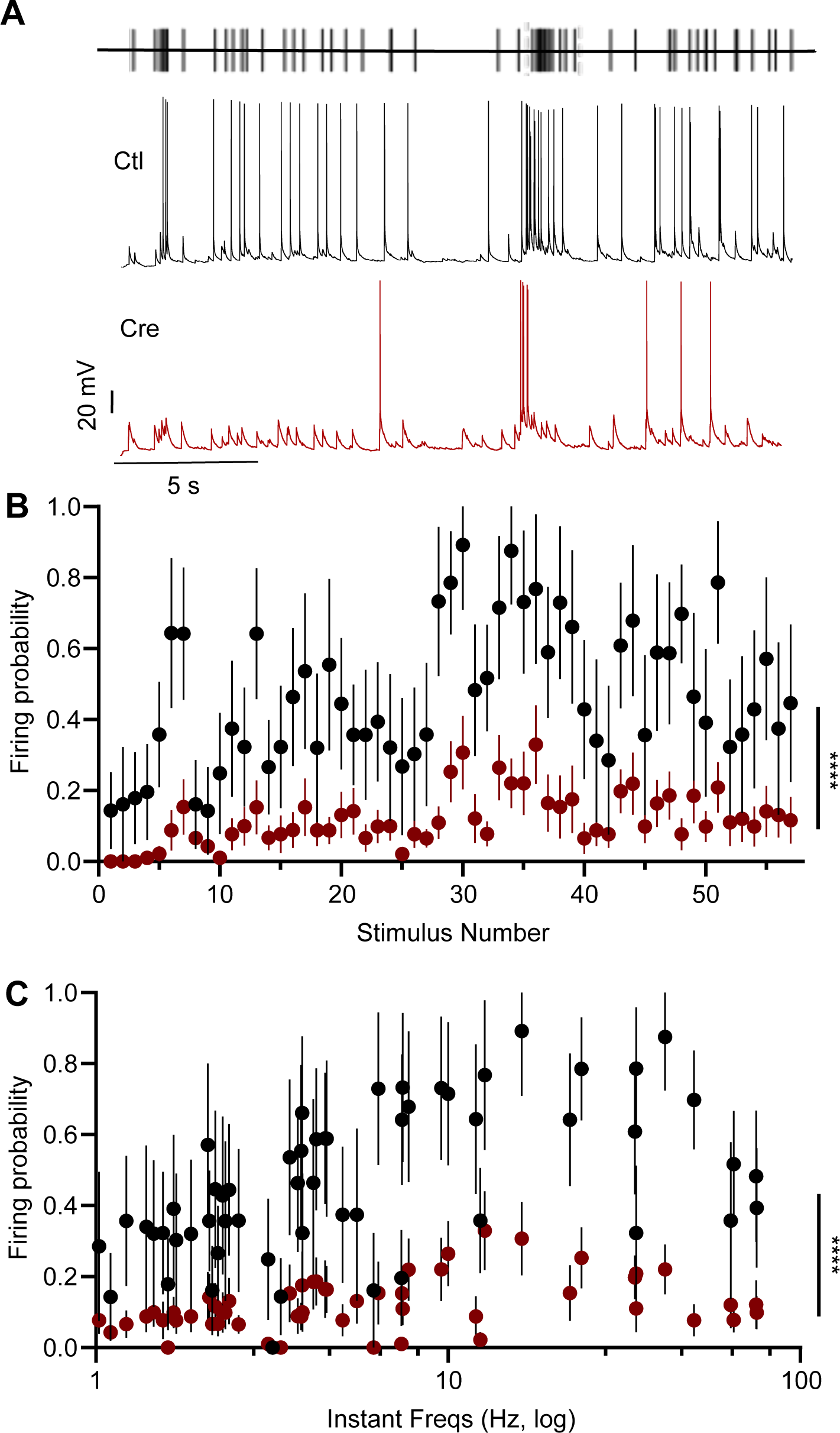
Abrogation of presynaptic short-term plasticity at Mf-CA3 synapses impairs spike transfer in response to natural pattern of stimulation. **A**. Schematic representation of the naturalistic stimulation pattern used to optically stimulate DG cells (top) while recording from CA3 PCs in the current clamp mode (bottom traces). Example traces of a single sequence of stimulation (20 s) for Ctl (black) and DG-Syt7 KO (red) mice. No drug was added. **B.** Graph of the mean firing probability for each of the 57 stimulations in the pattern, upon 7 repetitions of the 30 s sequence of stimulation pattern for Ctl (black) and DG-Syt7 KO (red) mice. **C.** Graph of firing probability in relation to instantaneous frequency of optical stimulation within the naturalistic pattern for Ctl (black) and DG-Syt7 KO (red) mice, showing a significant difference at all frequencies from 1 Hz to 75 Hz.

### *In vivo* electrophysiological recordings of DG-Syt7 KO mice: LFP in DG and CA3

With the previous slice experiments, we have validated a mouse model in which short-term presynaptic facilitation is selectively abrogated at Mf-CA3 synapses. To study the implications of this result in an *in vivo* context, we performed silicon probe recordings of ensemble neural activity in the dorsal DG and CA3 in head-fixed awake mice. We recorded from control and DG-Syt7 KO mice injected unilaterally. We verified post-hoc the viral infection of DG cells for LV-Control (n= 6) or LV-Cre (n = 9) mice (Fig 6A). We trained the mice to be maintained fixed with head bars while freely running on a wheel. Once the mice were habituated and showed no signs of stress, we performed 30 minute-long acute recordings in head-fixed mice using a 32-channel linear silicon probe lowered into the DG, and a 64-channel, 4-shank probe positioned in CA3 (Fig. 6A, B).

**Figure 6.**
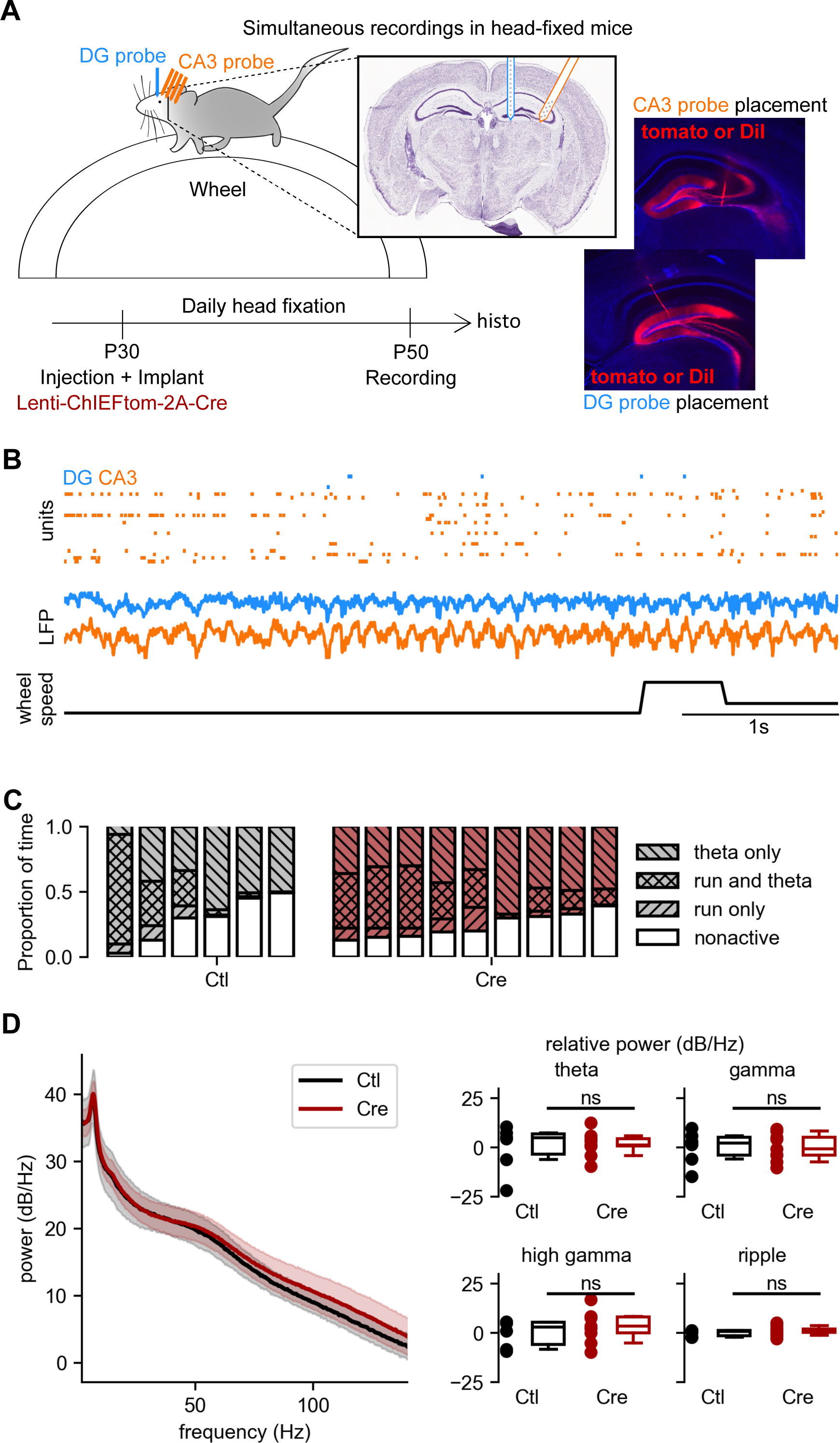
In vivo electrophysiological recordings in head-fixed awake mice: LFP in DG and CA3 **A.** *In vivo* recording setup. Mice were head-fixed and allowed to run freely on a wheel while silicon probe recordings were made in DG/CA1 and CA3. At around P30, mice were injected with either the Ctl or Cre viral constructs in the right dorsal hippocampus After recovery, they were habituated daily to head-fixation until the recording took place at around P50. **B.** Example recording. Top: raster plot of cell firing in DG and CA3. Middle: LFP from DG and CA3. Bottom: speed of the wheel during running. Scale bar is 1 second. **C.** Proportion of time in which theta and running were detected in the CA3 LFP and wheel speed, respectively. The union of theta and running were considered the ‘active’ state and all the rest was considered the ‘nonactive’ state. **D.** CA3 oscillations were not different between DG- Syt7 KO and control mice. Left: Average spectral density from DG-Syt7 KO and control mice (shaded regions represent standard deviation). Right: Relative power, defined as the difference from the mean of the control mice, was not different between Ctl and DG-Syt7 KO (Cre) mice for theta (4-12 Hz) (4.9 ± 7.8 dB/Hz [n = 6 mice] for Ctl, 1.2 ± 4.2 dB/Hz [n = 9 mice] for Cre, Mann Whitney, p = 0.78), gamma (20-50 Hz) (2.2 ± 6.2 dB/Hz [n = 6 mice] for Ctl, -0.6 ± 5.8 dB/Hz [n = 9 mice] for Cre, Mann Whitney, p = 0.95), high gamma (50-100 Hz) (3.0 ± 5.5 dB/Hz [n = 6 mice] for Ctl, 3.5 ± 6.0 dB/Hz [n = 9 mice] for Cre), and ripple frequency (100-300 Hz) (0.8 ± 1.2 dB/Hz [n = 6 mice] for Ctl, 1.7 ± 1.7 dB/Hz [n = 9 mice] for Cre, Mann Whitney, p = 0.46).

We analysed LFP recordings in CA3, and compared the proportion of time where theta and running were detected, the union of theta and running being considered the ‘active’ state and all the rest was considered the ‘nonactive’ state. Overall, no significant difference was detected between control and DG-Syt7 KO mice in the overall time spent in an active vs. inactive state (Fig. 6C). The power of oscillations in the delta (0.5-3.5 Hz), theta (4-12 Hz), gamma (20-50 Hz), high gamma (50-100 Hz) and ripple (100-300 Hz) frequency ranges between the two groups (Fig 6D) was further analyzed in both the DG and CA3. No difference was observed in the relative power in any of the frequency bands when comparing control and DG-Syt7 KO animals (Fig 6D). We next investigated whether the abrogation of presynaptic short-term facilitation at Mf-CA3 synapses could affect the LFP coherence between the DG and CA3. Using Welch’s method to calculate coherence, no statistically significant variation could be observed comparing the two groups (Sup Fig. 3A).

### *In vivo* electrophysiological analysis of spiking activity in DG and CA3

We then isolated single units in CA3 and compared firing frequencies for putative principal cells and interneurons (INs). Cell type classification was performed based on the waveform features: trough-to-peak, peak spectral power, and average firing rate were used to classify between putative INs and principal cells. The average number of recorded units was not different between Ctl and DG-Syt7 KO mice for CA3 PCs (Ctl = 55 ± 14, DG-Syt7 KO = 57 ± 14, Mann Whitney p = 0.9) and for CA3 INs (Ctl = 7 ± 3, DG-Syt7 KO = 7 ± 3, Mann Whitney, p = 0.8). Both Ctl and DG-Syt7 KO mice had smaller firing rates in CA3 PCs during the active state, but there was no difference between the two strains (Fig 7A) (bootstrap paired t test: Ctl, nonactive vs active p < 0.0001, DG-Syt7 KO, nonactive vs active p < 0.01, Mann Whitney: nonactive Ctl vs DG-Syt7 KO, p = 0.53, active Ctl vs DG-Syt7 KO, p = 0.61). CA3 INs had no strain-dependent or state-dependent differences in firing rate (Fig 7A) (bootstrap paired t test: Ctl nonactive vs active p = 0.19, DG-Syt7 KO nonactive vs active p = 0.9, Mann Whitney: nonactive Ctl vs DG-Syt7 KO, p = 0.95, active Ctl vs DG-Syt7 KO, p = 0.39).

**Figure 7.**
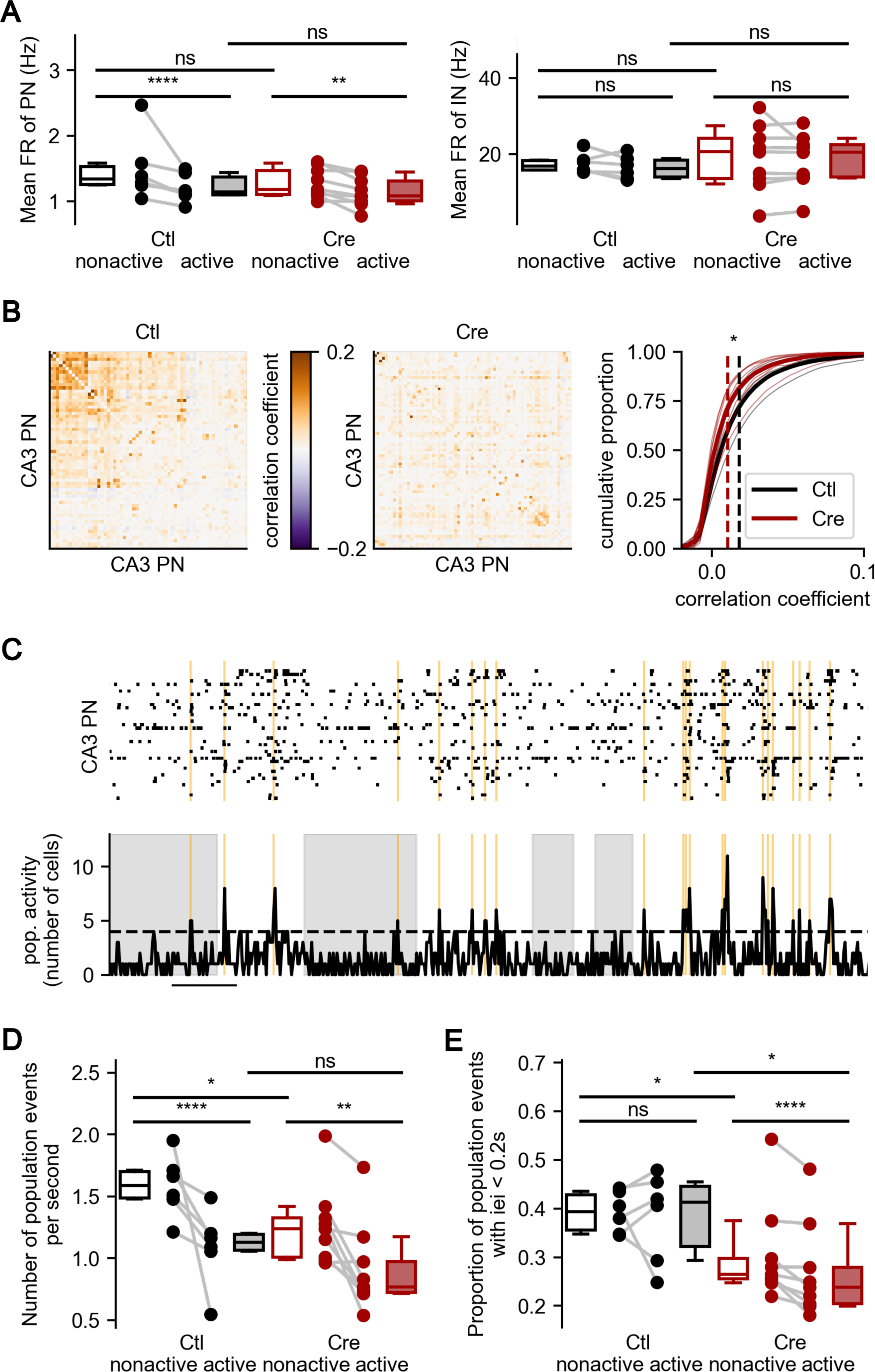
Reduction of CA3 pyramidal cell cofiring in DG-Syt7 KO mice, despite lack of changes in firing rate. **A.** State-dependent firing rates of CA3 pyramidal neurons (left) and interneurons (right). Both Ctl and DG-Syt7 KO (Cre) mice have smaller firing rates in CA3 PCs during the active state, but there is no difference between the two genetic conditions (1.3 ± 0.3 Hz [n = 6 mice] for Ctl PN nonactive state, 1.1 ± 0.2 Hz [n = 6 mice] for Ctl PN active state, 1.2 ± 0.2 Hz [n = 9 mice] for Cre PN nonactive state, 1.1 ± 0.2 Hz [n = 9 mice] for Cre PN active state, bootstrap paired t test: control nonactive vs active p < 0.0001, Cre nonactive vs active p < 0.01, Mann Whitney: nonactive control vs Cre p = 0.53, active control vs Cre p = 0.61). CA3 interneurons have no strain-dependent or state-dependent differences in firing rate (16.9 ± 2.0 Hz [n = 6 mice] for Ctl IN nonactive state, 16.1 ± 2.5 Hz [n = 6 mice] for Ctl IN active state, 20.6 ± 6.8 Hz [n = 9 mice] for Cre IN nonactive state, 20.4 ± 5.6 Hz [n = 9 mice] for Cre IN active state, bootstrap paired t test: control nonactive vs active p = 0.19, Cre nonactive vs active p = 0.9, Mann Whitney: nonactive control vs Cre p = 0.95, active control vs Cre p = 0.39). Data are expressed as median ± average absolute deviation from the median (MAD) here and in the rest of this figure. B. Correlation coefficients between pairs of CA3 pyramidal neurons. Correlation coefficient matrices for example control (left) and DG-Syt7 KO mice (centre). Right: cumulative distribution of the pairwise correlation coefficients, showing a significant decrease in DG-Syt7 KO mice (0.018 ± 0.003 [n = 6 mice] for Ctl, 0.010 ± 0.003 [n = 9 mice] for Cre, bootstrap t test, p = 0.045). **C.** Example raster plot of CA3 pyramidal neurons (top) and population activity (bottom) from a control mouse. Grey shading shows when the mouse is in the active state. The horizontal dotted line is the threshold for finding population events, which are shown in orange. Population events are defined as periods when more cells are firing in a 20 ms window than would be expected by chance. The scale bar represents 1s. **D.** Rate of occurrence of population events. Both DG-Syt7 KO mice and controls have fewer population events per second during the active state than the nonactive state. In comparison with controls, DG- Syt7 KO mice have a reduced rate of population events during the nonactive state (1.6 ± 0.2 Hz [n = 6 mice] for Ctl nonactive state, 1.1 ± 0.2 Hz [n = 6 mice] for Ctl active state, 1.2 ± 0.2 Hz [n = 9 mice] for Cre nonactive state, 0.8 ± 0.2 Hz [n = 9 mice] for Cre active state, bootstrap paired t test: control nonactive vs active p < 0.0001, Cre nonactive vs active p < 0.01, Mann Whitney: nonactive control vs Cre p = 0.05, active control vs Cre p = 0.22). **E.** Proportion of population events that have an inter event interval shorter than 200 ms. In comparison with controls, population events in DG-Syt7 KO mice are less likely to occur close together in time during both the nonactive and active states (0.39 ± 0.04 [n = 6 mice] for Ctl nonactive state, 0.41 ± 0.07 Hz [n = 6 mice] for Ctl active state, 0.26 ± 0.06 Hz [n = 9 mice] for Cre nonactive state, 0.24 ± 0.06 Hz [n = 9 mice] for Cre active state bootstrap paired t test: control nonactive vs active p = 0.81, Cre nonactive vs active p < 0.0001, Mann Whitney: nonactive control vs Cre p = 0.03, active control vs Cre p = 0.05).

To evaluate the dynamics of CA3 co-firing, we calculated the correlation coefficient of the spike trains of pairs of CA3 PCs (Fig 7B). When averaged over all cell pairs, the correlation coefficient showed a decrease in DG-Syt7 KO mice compared to Ctl mice (Fig 7B, bootstrap t test, p = 0.045). We then performed a population analysis to compare more generally the co-activity of CA3 PCs in Ctl vs. DG- Syt7 KO mice. Population events were defined as periods when more cells were firing in a 20 ms window than would be expected by chance in a shuffled dataset (Fig. 7C). The rate of occurrence of population events (Fig. 7D) was strongly decreased between the non-active and the active states, in both Ctl and DG-Syt7 KO mice (Ctl, p <0.0001; Cre, p <0.01). During the non-active state, but not during the active state, the rate of occurrence of population events was significantly reduced in DG-Syt7 KO mice as compared to Ctl mice (non-active state, p = 0.05; active state, p = 0.22, Mann Whitney). We next analysed the distribution of population events across state and genetic conditions, by measuring the proportion of population events that have an inter-event interval shorter than 200 ms (Fig. 7E). The proportion of population events occurring during this time window was not different between the non-active and active state in Ctl mice (p = 0.83), but was largely reduced in DG-Syt7 KO mice (p < 0.001). Overall, population events were less likely to occur close together in time (during both the nonactive and active states) in DG-Syt7 KO mice as compared to Ctl mice (non-active state, p = 0.03; active state, p = 0.05, Mann Whitney). Taken together, these findings suggest that DG-specific Syt7 deletion does not affect the firing rates of individual cells or the LFP in CA3, but impairs the ability of DG to control the co-activation of CA3 ensembles.

### Presynaptic short-term plasticity at Mf-CA3 synapses is dispensable to perform pattern separation tasks

The DG has been proposed to perform pattern separation, the ability to discriminate between overlapping input patterns before encoding (Hainmueller and Bartos, 2020; Kesner and Rolls, 2015). To interrogate the behavioral relevance of presynaptic short-term plasticity at Mf-CA3 synapses, we first interrogated the consequences of DG-selective deletion of Syt7 on the ability of mice in two pattern separation tasks. We first performed a spatial separation task in which a mouse was placed in a 1 x 1 m arena and allowed to explore two objects for 12 minutes. After an inter-trial interval of 30 minutes, the mouse was placed again in the field; one of the objects did not move but the other object was displaced by 20, 35 or 45 cm away from the fixed object in a random order (Fig. 8A). The interaction ratio, the percentage of time spent by the mouse exploring the displaced object over total time exploring both objects, is considered as a proxy for the spatial ability of identifying the object displacement. None of the mice interacted longer with the object moved by 20 cm. However, the mice significantly increased exploration of the object when displaced by 35 cm and 45 cm, suggesting that they were able to perform the pattern separation task, and no difference was observed between the control mice and DG-Syt7 KO mice (Fig. 8B).

**Figure 8.**
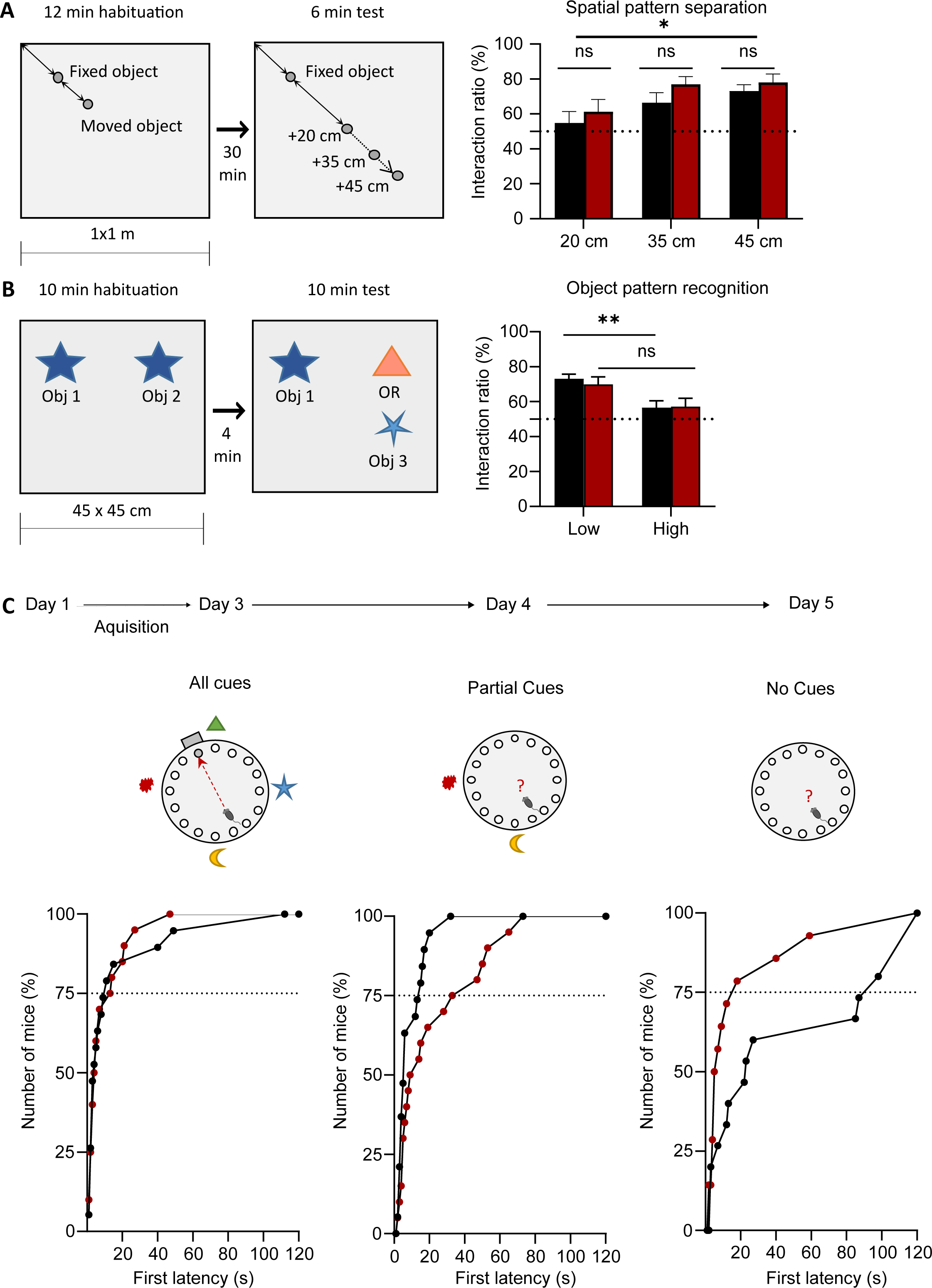
Abrogation of short-term plasticity at Mf-CA3 synapses does not affect pattern separation but impairs pattern completion. **A**. Abrogation of presynaptic facilitation at Mf-CA3 synapses does not impair the performance of mice in a spatial pattern separation (SPS) task. SPS results show no difference between Ctl and Cre injected mice when the moved object is displaced at 20, 35 and 45 cm of distance from the fixed object (at 20 cm: n = 20 Ctl and n= 12 Cre; at 35 cm: n = 20 Ctl and n = 13 in Cre; at 45 cm: n=20 Ctl, n= 13 Cre). **B**. Scheme for the description of an object pattern separation (OPS) task. OPS results (Ctl low = 73.2%, n=23, Cre low =70.0%, n=12; Ctl high = 56.6%, n=21, Cre high = 57.2%, n=12). The total interaction time, although not different for low similarity objects, is higher in DG-Syt7 KO mice for the high similarity objects (Two-way Anova, low p=0.1913, high p=0.0033).

Complementarily, we used a task based on visual separation of objects with different values of similarity. For this, we placed the mice in a 45 X 45 cm arena with two objects, and after an inter-trial period of 4 minutes, the mice were placed again in the arena with one of the two objects replaced by an object of either low or high similarity. The interaction ratio was calculated as the interaction time with the new object as compared to the total time exploring both objects (Fig. 8B). With high similarity objects, the ratio remained around chance levels, suggesting that the mice did no perceive the difference with the new object. In contrast, with low similarity objects, both control and DG-Syt7 KO mice significantly increased exploration of the new object to the same level, showing similar ability to perform the task. We observed however a tendency that the total interaction time with the objects to be higher in DG-Syt7 KO mice. Overall, these results indicate that the absence of presynaptic plasticity at Mf-CA3 synapses is dispensable to perform pattern separation in these spatial and object discrimination tasks.

### Abrogation of short-term presynaptic plasticity at Mf-CA3 synapses impairs pattern completion

CA3 has been proposed to be key for pattern completion, the ability to retrieve a full pattern of activity from partial or degraded inputs, based on the dense network of recurrent collaterals within CA3 PCs (Kesner and Rolls, 2015). To evaluate whether the abrogation of short-term plasticity at Mf-CA3 synapses impacted pattern completion, we implemented a spatial memory task in the Barnes maze, consisting of a large table with 18 holes, one of which is connected to an “escape box” allowing the mouse to hide from light and open space. During a three-day acquisition phase, the mice were placed in the Barnes maze with a complete set of external cues allowing the mice to rapidly learn to find the escape box (Fig. 8C). The DG-Syt7 KO mice traveled a longer latency to find the escape box on the first day as compared to Ctl mice (first latency = 70.0 sec in DG-Syt7 KO, first latency = 55.3 sec in Ctl, p = 0.0143, Two-way Anova). They however performed equally well as the control mice on the second and third day to find the escape box (Day 2 : first latency = 33.7 sec in DG-Syt7 KO, first latency = 20.3 sec in Ctl, p = 0.9997, Two-way Anova. Day 3 : first latency = 18.8 sec in DG-Syt7 KO, first latency = 55.0 sec in Ctl, p = 0.9861, Two-way Anova). On the following days, the ability of mice to retrieve the escape box location was assessed while some of the cues were removed (Fig. 8C). We measured the latency of the mice to first reach the hole target over 30 sec. When all cues were present >75% of the mice in both groups found the target area within 10 sec (Fig. 8C). With partial cues, DG-Syt7 KO mice took longer than the control mice to find the target area: the first latency for 75% of mice in the Ctl group was 11 sec, against 33 sec from DG-Syt7 KO mice. Finally, when all the cues were removed from the arena (minimal cues), DG-Syt7 KO mice further increased their time to reach the target area as compared to control mice: the first latency for 75 % of the Ctl group was 15.5 sec against 93 sec for DG-Syt7 KO mice. Overall these experiments indicate that abrogation of presynaptic short-term plasticity at Mf-CA3 synapses induce a pronounced deficit in pattern completion.

### Emotional processes are altered in DG-Syt7 KO mice

As indicated, the spread of viral infection was not limited to the dorsal DG (Sup. Fig. 1). We therefore tested whether emotional processes which depend on the ventral hippocampus (Fanselow and Dong, 2010) could be affected. For this we first analyzed locomotor activity of mice in an open field to test for passive anxiety. No difference was observed in the total distance, in the time spent in the center or in thygmotaxis between DG-Syt7 KO mice and control animals (Fig. 9A). However, in the elevated plus maze, DG-Syt7 KO mice spent proportionally less time in the open arms than Ctl mice, although the total distance travelled was not different between the two groups (Fig. 9B). We finally used a test for burying behaviors, which reflects active anxiety and obsessive – compulsive-like behavior. In this test, 20 beads were placed on top of the litter and the isolated mice were placed in the cage for 20 minutes. The burying rate was calculated as the percentage of bead surface visible from the top of the cage after and before placing the mouse in the cage. Strikingly, DG-Syt7 KO mice exhibited much more burying than the control mice (Fig. 9C). Overall these experiments suggest that abrogation of presynaptic short-term facilitation at Mf-CA3 synapses in mice increased anxiety and promoted an obsessive – compulsive-like behavior, processes which may rely on the ventral hippocampus.

**Figure 9.**
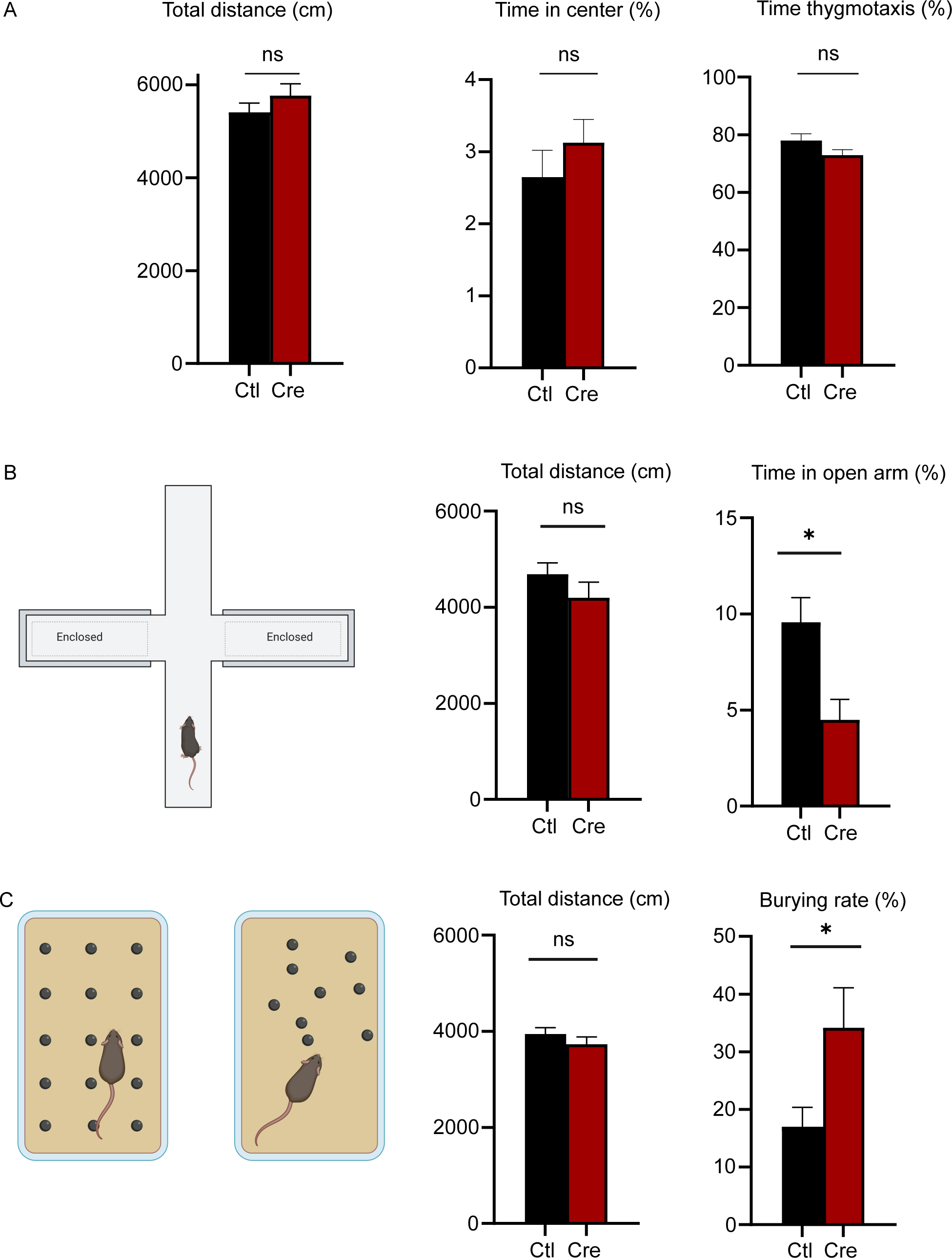
Abrogation of presynaptic facilitation at Mf-CA3 synapses impairs emotional processes. **A**. Control and Cre mice do not exhibit different behaviors in an open field (Ctl n = 23, Cre n = 12). Total distance is the same (5403 cm for Ctl, 5762 cm for Cre, unpaired t-test, p=0.2874), the percentage of time spent in the center is not changed (Ctl = 2.65 %, Cre = 3.13%, unpaired t-test, p = 0.4046), and the percentage of time expressing thygmotaxis is the same (Ctl = 78.0%, Cre = 73.0%, unpaired t-test, p=0.1668). **B**. After being exposed for 10 minutes in an elevated plus maze, although the total distance is the same within the two groups (Ctl = 4684 cm, n=23; Cre = 4198 cm, n =11, unpaired t-test, p = 0.2442), the time spent in the open arm is significantly lower in Cre mice (Ctl = 9.57%, Cre = 4.49%, unpaired t-test, p = 0.0168). **C**. After 15 minutes in a cage with bedding with 20 marbles, although the total distance roamed in the cage is the same (Ctl = 3944 cm, n = 23; Cre = 3729 cm, n = 13, unpaired t-test, p = 0.3203), the burying rate was significantly greater in Cre injected mice (Ctl = 17%, Cre = 34.14%, unpaired t-test, p = 0.0168).

## DISCUSSION

Synapses undergo plastic changes over a wide range of time windows, from ms to hours or days. For decades, a large body of work has addressed the implication of long-term synaptic plasticity (lasting >tens of minutes), mostly expressed at a postsynaptic level, in memory (Takeuchi et al., 2014). At most synapses in the brain, short-term plasticity (lasting from ms to minutes) dynamically modulates synaptic strength (Zucker and Regehr, 2002). Much less is known about the role of short term plasticity in the activity of synaptic circuits in relation to memory formation, such as in the hippocampus (Jackman and Regehr, 2017; Monday et al., 2018). Here we have validated and used a mouse model which displays selective abrogation of short-term facilitation at Mf-CA3 synapses, and we have examined the consequences on neural ensemble activity in the DG and CA3 as well as in memory formation.

*Validation of a mouse model with abrogation of presynaptic facilitation at Mf-CA3 synapses* Presynaptic facilitation is a form of short-term synaptic plasticity which can lead to a major increase in the strength of synaptic transmission at certain types of synapses, notably at synapses between layer 6 cortical pyramidal cells and thalamic relay cells (Deschênes & Hu, 1990) or at Mf-CA3 synapses (Nicoll and Schmitz, 2005). The large dynamic range of presynaptic facilitation at DG-CA3 connections synapses endows Mf-CA3 synapses with “conditional detonator” properties (Henze et al., 2002; Sachidhanandam et al., 2009; Vyleta et al., 2016), thought to be critical for directing the storage of information in the auto-associative CA3 network (Kesner and Rolls, 2015). This implies that activation of a single mossy fiber is sufficient to reliably discharge a postsynaptic CA3 PC following short bursts of incoming action potentials from the DG (Henze et al., 2002).

In order to selectively address the role of presynaptic facilitation at Mf-CA3 synapses (their “detonator properties”) on the activity of CA3 circuits, we took advantage of the fact that genetic deletion of Syt7 in constitutive KO mice strongly reduces presynaptic facilitation induced by 20 Hz trains stimulations at Mf-CA3 synapses (Jackman et al., 2016). We used Syt7 Floxed mice and DG selective viral tools to remove Syt7 from presynaptic Mf terminals in CA3. Post-hoc immunohistological examination, indicated that Syt7 was decreased by about 60% upon infection of the dorsal DG with the LV-Cre construct, which likely pertains to an equivalent percentage of infected DG GCs. Presynaptic facilitation in response to 20 Hz trains of 10 stimulations was almost entirely abrogated following the deletion of Syt7 at Mf-CA3 synapses, from a facilitating factor of around 6 in control mice to a factor of 1.5 in DG- Syt7 KO mice. The remaining facilitation, which was small, may rely on the broadening of presynaptic K^+^ channels following trains of incoming action potentials, which translates into higher Ca^2+^ influx and facilitated release (Geiger & Jonas, 2000). Our detailed analysis of Mf-CA3 synapses in slices from DG- Syt7 KO mice provided a clear indication that the basal synaptic properties of Mf-CA3 synapses (quantal content, probability of release, number of active release sites) are likely unchanged. In order to establish a reliable correlation between presynaptic short-term facilitation, *in vivo* hippocampal electrophysiological activity and behavior observed in these mice, it was important to pinpoint whether other plasticity mechanisms known to occur at Mf-CA3 synapses were also impaired.

In particular, Mf-CA3 synapses display both presynaptic PTP and presynaptic LTP, which can be triggered by a short series of high frequency trains (Salin et al., 1996). Whereas evoked Mf-CA3 synaptic responses during the induction trains obtained by optical stimulation were drastically reduced in DG-Syt7 KO mice, the extent and time course of both PTP and LTP were not significantly altered. This strongly suggests that synaptic release is not required for the expression of PTP and LTP. Hence, both PTP and LTP which require the activation of PKA (Monday et al., 2018; Vandael & Jonas, 2024; WeissKOpf et al., 1994) (Marneffe et al, accepted for publication), may be triggered by cell-autonomous increase of cAMP in the presynaptic Mf terminal upon the incoming train of action potentials. Therefore, it seems that Syt7 is dispensable for the expression of PTP and LTP. As a consequence, the DG-Syt7 KO mice selectively harbor defective short-term facilitation on the ms- second time scale.

Monosynaptic innervation from Mfs to GABAergic CA3 interneurons creates the anatomical basis for robust feedforward inhibition of CA3 PCs (Rebola et al., 2017). The impact of DG selective deletion of Syt7 is likely to be different dependent on the synapses with distinct types of GABAergic interneurons in CA3 contacted by Mfs, as these synaptic connections display cell type-specific properties and short-term plasticity rules (Acsády et al., 1998; Szabadics & Soltesz, 2009). It is important however to understand how, overall, suppression of Syt7 at Mf-CA3 synapses impacts spike transfer between the DG and CA3. We have taken a step towards elucidating this by comparing the extent and short-term plasticity of feedforward inhibition at Mf-CA3 synapses in control and DG-Syt7 KO mice. Although the amplitude of disynaptic IPSCs triggered by the optical stimulation of DG granule cells showed high variability, the amplitude was overall reduced at all stimulation frequencies tested (0.1 Hz, 1 Hz and 20 Hz), indicating less efficient feedforward inhibition. However, because IPSCs on average do not facilitate in response to a train of Mf stimulation (also see (Zucca et al., 2017)), the excitation/inhibition ratio was strongly dependent on the facilitation of EPSCs. Hence the increase of the E/I ratio during a train of stimulation was not present in DG-Syt7 KO mice. Future work should directly analyse short-term plasticity at connections between Mfs and CA3 INs in DG-Syt7 KO mice, taking into account their large diversity (Szabadics & Soltesz, 2009). Overall, the loss of presynaptic facilitation at Mf-CA3 PC synapses is, at least in part, counterbalanced by a reduced feedforward inhibition. It is interesting to note that reduced feedforward inhibition may be viewed as a compensatory homeostatic consequence of the suppression of facilitation at Mf-CA3 PC synapses. Structural plasticity of Mf-CA3 connections onto CA3 interneurons, through changes in the number of filopodia, may be involved, as it provides a key mechanism for regulating the interface between the DG and CA3, e.g. following one-trial learning (Ruediger et al., 2011). Nevertheless the reduced feedforward inhibition in DG-Syt7 KO mice did not prevent a major failure in spike transmission from DG to CA3, in conditions of natural spiking activity.

### Hippocampal activity in the absence of the detonator properties at Mf-CA3 synapses

Our subsequent analysis provides a mechanistic link between the dynamic properties of short-term plasticity at Mf-CA3 synapses and hippocampus-wide networks *in vivo*. Due to the high dynamic range of presynaptic plasticity, Mf-CA3 synapses are “conditional detonators”, allowing a single DG granule cell to reliably discharge a CA3 PC depending on the granule cell firing pattern (Henze et al., 2002). *In vivo*, patch-clamp recordings of CA3 PCs combined with optogenetic activation of DG cells have shown that the tight balance between excitation and inhibition at Mf-CA3 synapses provides a sharp window of input frequencies (around 10 Hz) at which spike transfer is maximized (Zucca et al., 2017). Similarly, coherent slow gamma between DG and CA3 has been shown to be associated with the encoding phase of a memory task (Trimper et al., 2017). With these features in mind, we analysed the correlation of LFP oscillations between the DG and CA3 in control and DG-Syt7 KO mice. Unexpectedly, we found no changes in the power of oscillations in all frequency bands from delta to sharp wave ripple (SWR) frequencies in both the DG and CA3, and the coherence between these oscillations across these two connected regions was not affected either. SWRs are prominent hippocampal network oscillations (150–250 Hz) observed during sleep as well as during periods of immobility. Their occurrence in CA3 is biased by inputs from the DG and entorhinal cortex (Sullivan et al., 2011). Furthermore, during a working memory task, Mf inputs to CA3 were shown to be required for the generation of SWRs (Sasaki et al., 2018). Therefore, a further analysis of the occurrence of SWRs in reference to DG activity is certainly necessary. It should also be stressed that the recordings were performed in awake mice allowed to freely run on a wheel, but which were not subjected to a specific task. We cannot exclude that differences in oscillatory activity in CA3 and in the cross-correlation between the DG and CA3 may be observed in behavioral conditions which increase the activity of the DG, such as in a memory task.

Whereas we found no difference in the general firing pattern of CA3 PCs (average frequency, bursting mode) and INs between control and DG-Syt7 KO mice, we observed a striking difference in the correlated spiking activity among CA3 PCs. Pairwise cross-correlations of CA3 PC spiking, as well as population activity, were disrupted in the absence of Mf-CA3 presynaptic facilitation. This suggests that CA3 PCs which fire together, possibly by virtue of their recurrent connectivity, may be controlled by common DG granule cells. The firing regime of these DG cells allows to facilitate downstream the spiking in a connected group of CA3 PCs through presynaptic facilitation of Mf-CA3 inputs. It is known that a single DG cell contacts up to 15 CA3 PCs (Rebola et al., 2017; Vandael & Jonas, 2024), but it is still unclear whether the CA3 PCs connected by a single DG cell are themselves part of an ensemble through reciprocal connections (Rebola et al., 2017). Two photon imaging of CA3 neurons over days has identified ensembles of co-active CA3 PCs which are stable across days (Schoenfeld et al., 2021), a feature which may participate in the stable encoding of spatial information (Stefanini et al., 2020). Overall, our data indicate that disarming the detonator Mf-CA3 synapses does not result in a total disconnection between the DG and CA3. However, from the knowledge gathered here, short term facilitation of Mf-CA3 synapses appears to be engaged in the controlled activity of connected ensembles of CA3 PCs. It remains to be demonstrated whether specific behavioral conditions and tasks would unmask larger differences in the activity CA3 PCs in the absence of a detonator input from the DG.

### Presynaptic facilitation is required for pattern completion, but not pattern separation

Within the hippocampus, the DG-CA3 circuit is thought to play a critical role in pattern separation during learning, a mechanism by which similar or overlapping inputs are made divergent at the level of output to CA3 (Kesner & Rolls, 2015; Neunuebel & Knierim, 2014). Rather unexpectedly, the abrogation of presynaptic plasticity at Mf-CA3 synapses did not lead to any impairment in the DG-Syt7 KO mice in two memory tasks which are thought to rely on pattern separation (Cès et al., 2018). This may be explained by the fact that young adult born DG neurons play an essential role in behavioral pattern separation (Danielson et al., 2016; Nakashiba et al., 2012), and that this process possibly requires Mf-CA3 short-term plasticity for efficient spike transfer between DG and CA3. At the moment, we have yet to assess whether neurons infected by the LV-Cre construct comprise adult born neurons, and if so, whether these neurons are already in a mature state.

In contrast to the DG, CA3 has been proposed to perform pattern completion, the ability to retrieve a full pattern of activity from partial or degraded inputs, based on the extensive recurrent collateral connections between CA3 PCs (Kesner & Rolls, 2015). We observed that DG-Syt7 KO mice performed equally well than control mice to find the escape box in the Barnes maze when full cues were present, but their ability in conditions of degraded cues was largely compromised, consistent with impaired pattern completion. As discussed above, a major feature impacted in DG-Syt7 KO mice is a reduced level of co-activity of CA3 PCs. We postulate that in the absence of efficient Mf-CA3 connections through presynaptic facilitation, CA3 PCs participating in a neural ensemble encoding for a spatial location are less capable of retrieving the full neural pattern because of decreased correlated activity, and possibly impaired synaptic plasticity between CA3 PCs. Synaptic plasticity within the CA3 recurrent network has been proposed to be important for memory retrieval (Nakazawa et al., 2002). On the other hand, Mf-CA3 synaptic inputs are involved in heterosynaptic plasticity; LTP at CA3–CA3 PC synapses can indeed be induced by the coincident firing of brief spike trains of Mfs and of associative/commissural inputs (KObayashi & Poo, 2004; Sachidhanandam et al., 2009). Hence, abrogation of presynaptic plasticity at Mf-CA3 synapses may impact on synaptic plasticity within CA3 engrams, and consequently impair the retrieval of memory through pattern completion.

### Abrogation of presynaptic plasticity impairs the emotional state

We found that DG-Syt7 KO mice were more anxious and expressed an obsessive–compulsive-like behavior. It is well known that the ventral hippocampus is a regulator of anxiety and emotional responses (Fanselow & Dong, 2010; Kheirbek et al., 2013). We did not specifically target the ventral hippocampus, but the LV-Cre injected in the dorsal hippocampus largely spread to the ventral part (see Sup Fig. 1). The morphological and functional features of the Mf track and Mf-CA3 synapses is conserved along the dorso-ventral axis (Li et al., 2024), including presynaptic facilitation. Interestingly, Syt7 was identified as a candidate risk factor for bipolar disorders in human, and the full Syt7 KO mice showed manic-like behavioral abnormalities in the dark phase and depressive-like behavioral abnormalities in the light phase (Shen et al., 2020). In the future, the possibility that these features rely at least in part on dysfunctional DG-CA3 connections should be examined.

Overall, the use of DG-Syt7 KO mice to selectively disarm the conditional detonator properties at Mf- CA3 synapses had a limited impact on the firing rates and LFP power in the DG and CA3, but strongly reduced the correlated activity of CA3 PCs within neural ensembles. These features appear to correlate with a lack of consequences of the genetic manipulation on behavioral tasks which require pattern separation, but may explain the impact on the retrieval of spatial memory through pattern completion. Future experiments are needed to further examine the impact of disarming the detonator synapse on the activity of neuronal ensembles in a mouse subjected to a memory task.

