## Supplementary Figures 1 to 3 for "Abrogation of presynaptic facilitation at hippocampal mossy fiber synapses impacts neural ensemble activity and spatial memory"

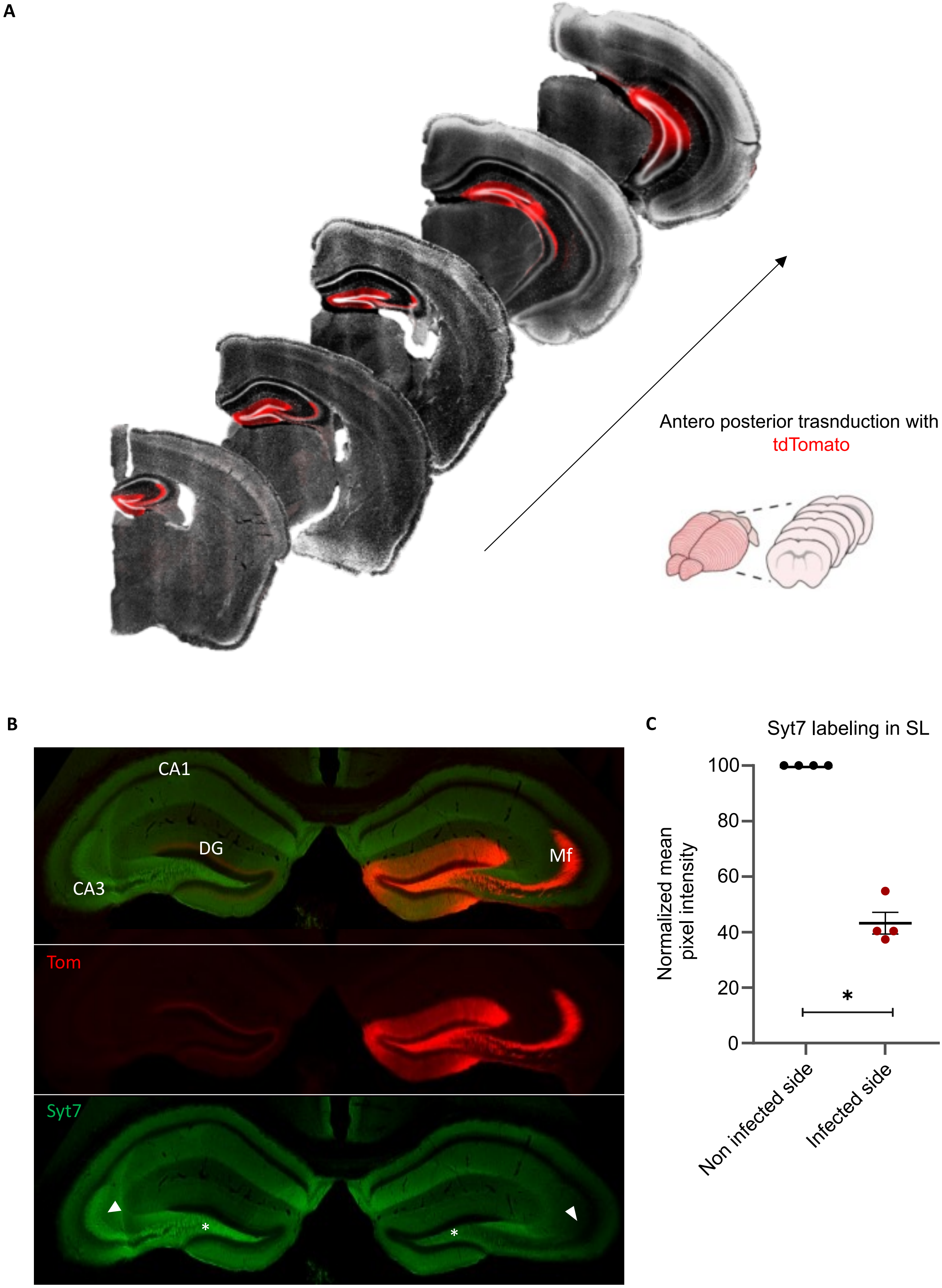


Supplementary figure 1

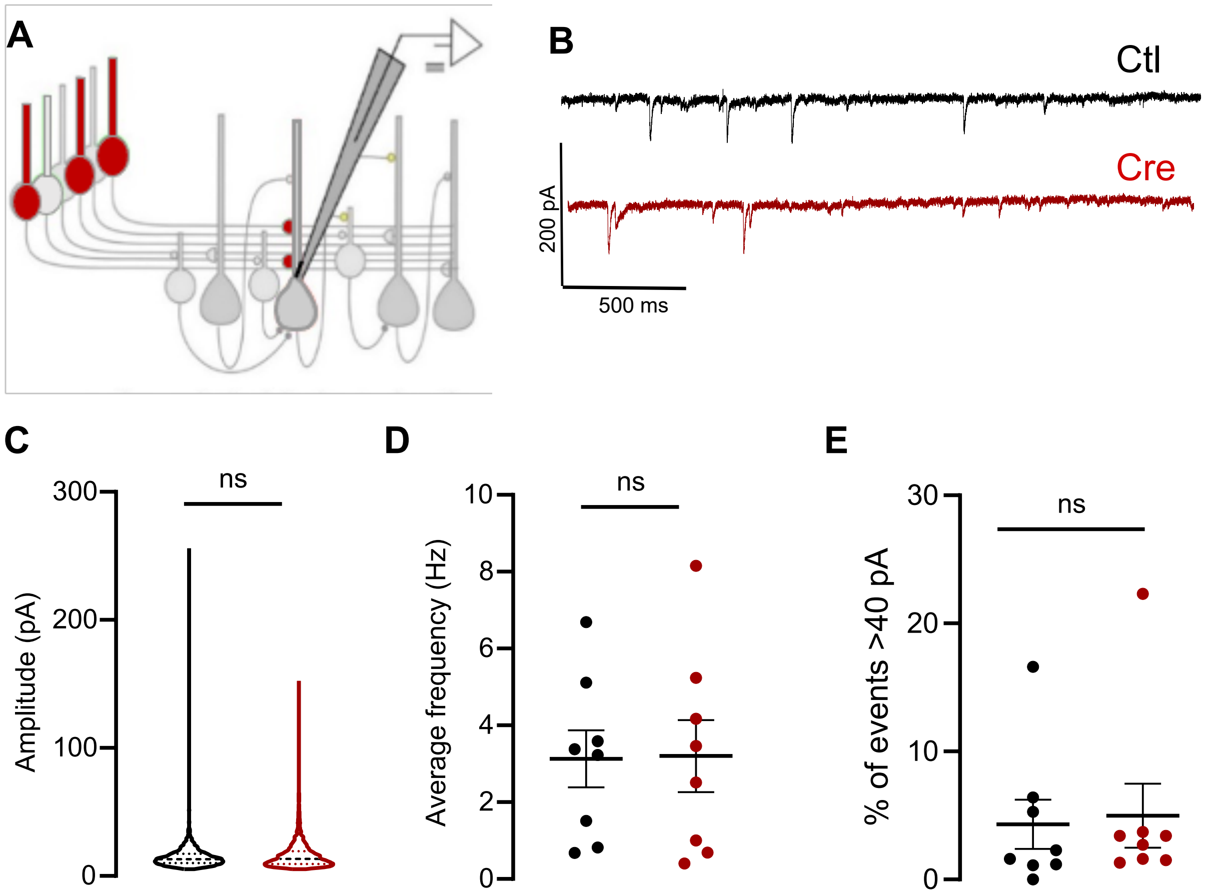


Supplementary figure 2

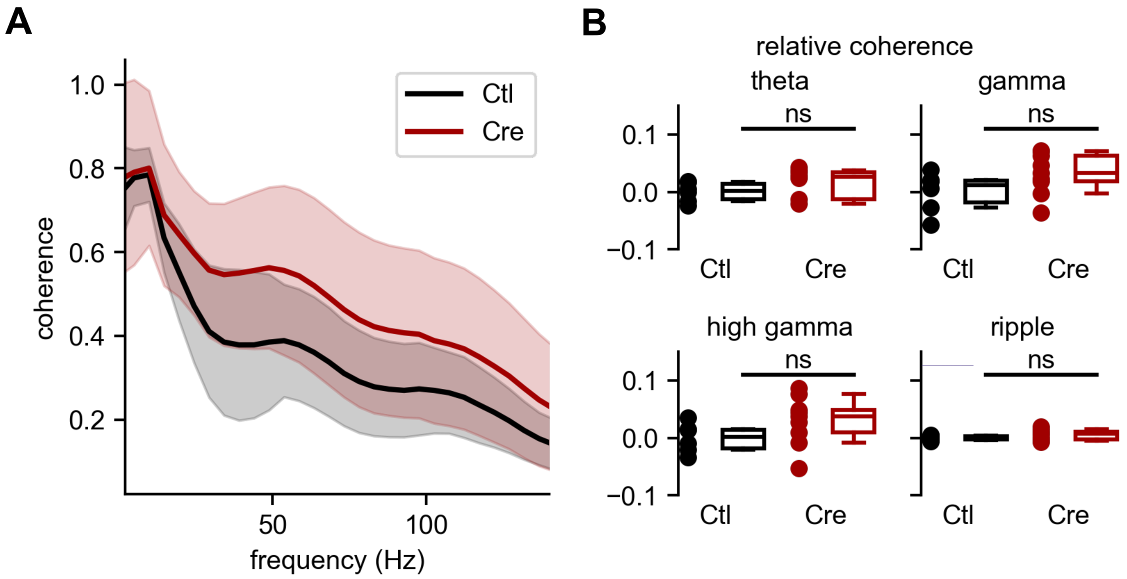


Supplementary figure 3

### Legends Supplementary figures

**Supplementary Figure 1 - Distribution of transduced DG neurons along the dorso-ventral axis**

Distribution of the transduction of DG neurons along the dorso-ventral axis, following injection of the LV-Cre construct in the dorsal hippocampus. The transduction of transduction speard from the dorsal to the ventral hippocampus, as indicated by td Tomato labelling of DG cells.

**Supplementary Figure 2 : Abrogation of short-term facilitation at Mf-CA3 synapses does not affect spontaneous synaptic activity in CA3 PCs.**

**A**. Patch clamp recordings performed in CA3 PCs in presence with 50 µM D-AP5. **B**. Example traces of spontaneous recordings in control and DG-Syt7 KO mice. **C**. Left panel represents the average amplitude of recorded spontaneous EPSCs (Mean amplitude = 14.8 pA in ctl over n=7868 events and mean amplitude = 16.4 pA over n= 8056 events in DG-Syt7 KO, Mann-Whitney test, p = 0.1917). **D**. Average frequency of spontaneous EPSCs (AF = 3.1 pA in ctl and AF = 3.2 pA in DG-Syt7 KO, Mann-Whitney test, p> 0.9). **E**. Proportion of EPSCs >40 pA, likely representing EPSCs of Mf origin in control vs. DG-Syt7 KO (n= 4.31 in Ctl and n = 4.98 in DG-Syt7 KO, Mann-Whitney test, p = 0.5908).

**Supplementary Figure 3. LFP coherence in DG-CA3 is not affected in DG-Syt7 KO mice.**

**A**. Average coherence between DG and CA3 LFP from DG-Syt7 KO and control mice (shaded regions represent standard deviation). **B.** Relative coherence, defined as the difference from the mean of the control mice, was not different between DG-Syt7 KO and control mice for theta (4-12 Hz) (0.002 ± 0.013 , n = 6 mice for Ctl, 0.027 ± 0.030, n = 9 mice for Cre, Mann Whitney, p = 0.18), gamma (20-50 Hz) (0.012 ± 0.026 [n = 6 mice] for Ctl, 0.033 ± 0.028 [n = 9 mice] for Cre, Mann Whitney, p = 0.14), high gamma (50-100 Hz) (0.002 ± 0.021, n = 6 mice for Ctl, 0.037 ± 0.031, n = 9 mice for Cre, Mann Whitney, p = 0.11), and ripple frequency (100-300 Hz) (0.000 ± 0.003, n = 6 mice for Ctl, 0.007 ± 0.007 [n = 9 mice] for Cre, Mann Whitney, p = 0.33).
